# A transient decrease in mitochondrial activity is required to establish the ganglion cell fate in retina adapted for high acuity vision

**DOI:** 10.1101/2020.03.23.002998

**Authors:** Laurent Brodier, Tania Rodrigues, Lidia Matter-Sadzinski, Jean-Marc Matter

## Abstract

Although the plan of the retina is well conserved in vertebrates, there are considerable variations in cell type diversity and number, as well as in the organization and properties of the tissue. The high ratios of retinal ganglion cells (RGCs) to cones in primate fovea and bird retinas favor neural circuits essential for high visual acuity and color vision. The role that cell metabolism could play in cell fate decision during embryonic development of the nervous system is still largely unknown. Here, we describe how subtle changes of mitochondrial activity along the pathway converting uncommitted progenitors into newborn RGCs increase the recruitment of RGC-fated progenitors. ATOH7, a proneural protein dedicated to the production of RGCs in vertebrates, activates transcription of the Hes5.3 gene in pre-committed progenitors. The HES5.3 protein, in turn, regulates a transient decrease in mitochondrial activity via the retinoic acid signaling pathway few hours before cell commitment. This metabolic shift lengthens the progression of the ultimate cell cycle and is a necessary step for upregulating Atoh7 and promoting RGC differentiation.

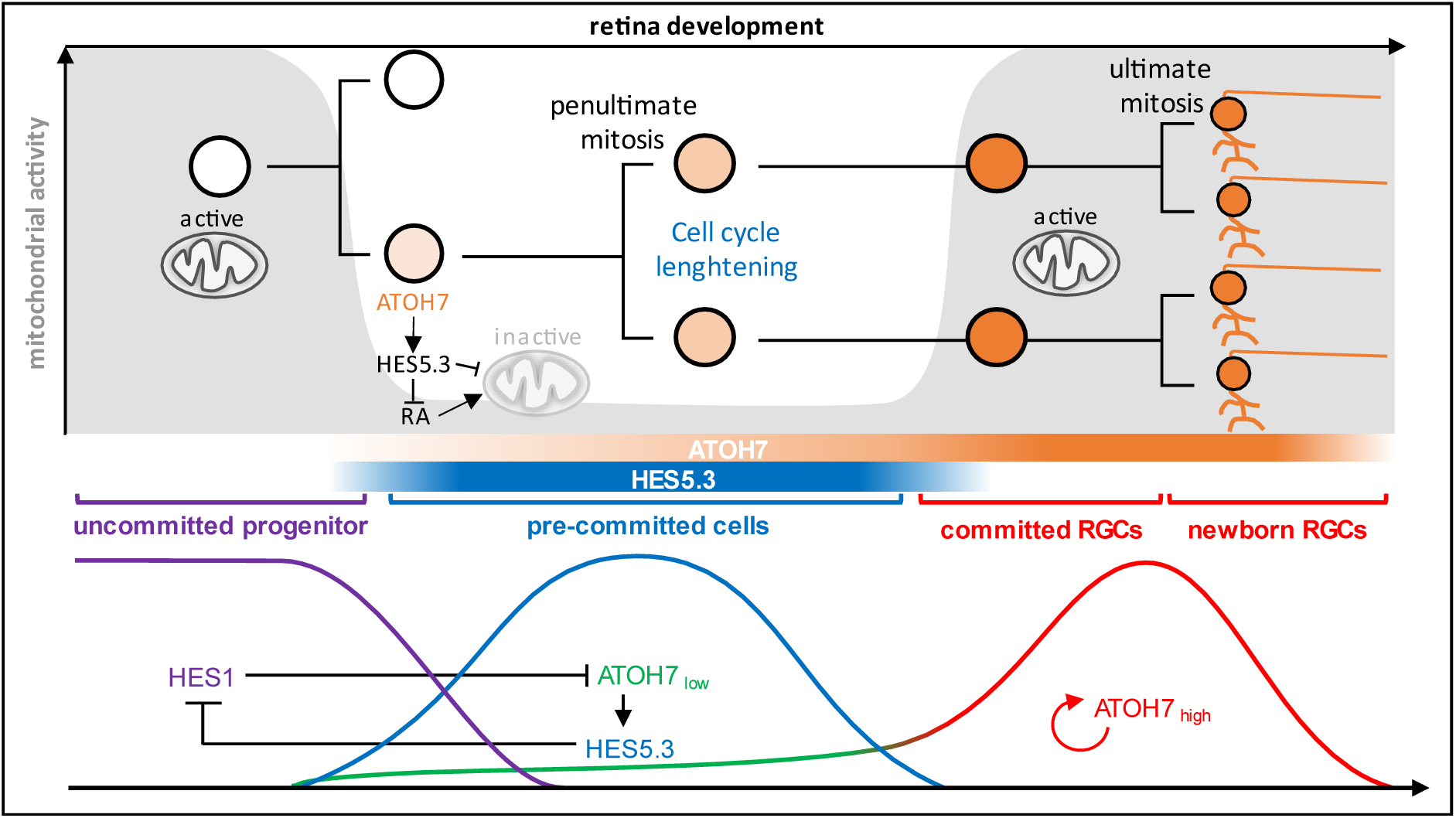
GRAPHICAL ABSTRACT.

## INTRODUCTION

Identifying a link between metabolic state and control of gene activity during cellular differentiation is a fascinating emerging field under intense investigation in different systems. Yet, little is known about the role of metabolic rewiring during development of the CNS. The main challenge lies in the tracing of metabolic activities with sufficient spatiotemporal resolution during neurogenesis and cell differentiation. The retina and particularly the retinal ganglion cells (RGCs) are among the largest ATP consumers in the whole body. Adult RGCs undergo continuous electrical activity and collectively transmit visual information to the brain. Several metabolic diseases have been linked to deficits in vision, and mitochondria are recognized as players in development of eye diseases like Leber’s hereditary optic neuropathy (Jarrett et al., 2010). Moreover, defective mitochondria are thought to participate in the degeneration of optic nerve axons in glaucoma (Osborne et al., 2006; Osborne et al., 2016; Tezel, 2006). RGC is a type of neuron located near the inner surface of the retina. It receives visual information from photoreceptors via two intermediate neuron types: bipolar cells and amacrine cells. RGC axons exit the retina through the optic disk, where they bundle together to form the optic nerve. RGC axons are unmyelinated before they exit the eye (Andrews et al., 1999; Bristow et al., 2002), a feature that decreases light scattering but which requires high and sustained energy supply for action potential propagation.

Mitochondria are central to energy production by oxidative phosphorylation (OXPHOS) (Rich and Marechal, 2010). While RGCs rely largely on OXPHOS for energy supply, recent experiments confirmed Warburg’s original findings suggesting the importance of aerobic glycolysis in photoreceptors (Chinchore et al., 2017; Ng et al., 2015; Warburg, 1925). Apart from their classical role as the “powerhouse of the cell”, mitochondria have been credited with many other regulatory functions (McBride et al., 2006). In particular, they could participate in cell fate decision and differentiation (Folmes et al., 2012). In Xenopus retina, progenitor cells are glycolytic, and the precocious differentiation of progenitors into RGCs is sufficient to induce a switch to OXPHOS (Agathocleous et al., 2012). A different situation occurs in the developing mouse retina where neuroblasts rely on OXPHOS for energy supply, whereas glycolysis and low mitochondrial content are required for RGC differentiation (Esteban-Martinez et al., 2017). Significant interspecies differences in metabolism could reflect variations in RGC number, RGC density and in the ratio of RGCs to photoreceptors. We might wonder to what extent conclusions reached in animal model that have no macula and no fovea are relevant to metabolic requirement of retinas developing these highly specialized regions. With a high ratio of RGCs to cones and a rod-free area, the avian retina resembles the primate macula and fovea (Da Silva and Cepko, 2017; Querubin et al., 2009; Rodrigues et al., 2016). This prompted us to investigate mitochondrial dynamics and activity along the pathway converting uncommitted progenitors into newborn RGCs in chick and pigeon retinas. We show that active mitochondria are abundant in uncommitted progenitors as well as in newborn RGCs. Atonal homolog 7 (ATOH7, also known as Ath5), is required for the production of RGCs in vertebrates (Brown et al., 2001; Del Bene et al., 2007; Kanekar et al., 1997; Kay et al., 2001; Liu et al., 2001; Matter-Sadzinski et al., 2001; Wang et al., 2001) and the *Hes5.3* gene (ENSGALG00000001136) is one of its earliest target. HES5.3 promotes, via cell cycle lengthening, the transition from pre-committed progenitors into cells committed to the RGC fate (Chiodini et al., 2013). Here we show that, while uncommitted progenitors have high contents of active mitochondria, mitochondrial activity drops in Hes5.3+ pre-committed cells. HES5.3 decreases mitochondrial activity by lowering retinoic acid (RA) level through the activation of Cyp26A1. Transient inactivation of mitochondria slows down the cell cycle progression, thereby enhancing the ability of ATOH7 to induce RGCs.

## RESULTS

### Apical accumulation of mitochondria at the onset of cell differentiation

To visualize mitochondria in vivo, we electroporated chick embryonic retinas with a plasmid expressing the DsRed2 fluorescent protein fused to the mitochondrial targeting sequence of human cytochrome c oxidase subunit VIII under the control of a constitutive CMV promoter (CMV-MitoDsRed2). A membrane potential is required to send proteins to the inner mitochondrial membrane (Hood et al., 2003; Rehling et al., 2001) and we assume that this is valid for MitoDsRed2. MitoTracker Green FM (MTG) is retained in mitochondria irrespectively of the loss of membrane potential (Elmore et al., 2004), and we found that only a subset of MTG-labelled mitochondria were labelled with MitoDsRed2 (Fig. S1 A-E). Likewise, the mitochondria membrane potential indicator tetramethylrhodamine methyl ester (TMRM) did not mark all MTG positive mitochondria, suggesting that a fraction of mitochondria have lower membrane potential. TMRM fluorescence disappeared 5 to 10 minutes after addition of 5 µM Carbonyl cyanide-4-(trifluoromethoxy)phenylhydrazone (FCCP), a mitochondrial uncoupler, whereas this inhibitor had no effect on the MTG signal (Fig. S1 F). Equivalent fractions of MTG positive mitochondria were labelled with MitoDsRed2 or TMRM, suggesting that MitoDsRed2 and TMRM label mitochondria with high membrane potential (Fig S1 E). First, we asked whether mitochondria displayed specific intracellular distribution along the apico-basal axis, which can be associated with specific cell cycle or cell differentiation phases. In chick retinas electroporated with CMV-MitoDsRed2 and CMV-GFP reporter plasmids at E5 and processed for confocal microscopy 24 h later, mitochondria preferentially accumulated on the apical side of the retina (Fig. 1A, D). To determine whether this apical concentration was a general feature of tissue growth or was related to cell differentiation, we compared the distribution of mitochondria in chick and pigeon retinas. Although retinas grow at a similar pace in both species, cell differentiation is delayed by ∼3 days in pigeon (Rodrigues et al., 2016). In pigeon and chick progenitors, mitochondria were distributed along the apico-basal axis (Fig. 1B, C), and the accumulation of fluorescent mitochondria on the apical side at E5 was more pronounced in chick than in pigeon retina (Fig. 1A, B, D). In order to ascertain that MitoDsRed2 labeling was reflecting mitochondria distribution, we analyzed chick and pigeon retinas by transmission electron microscopy (TEM). Mitochondria densities were quantified on the apical and basal sides as the fraction of cytoplasm occupied by the organelles. While, in chick, mitochondria were already concentrated apically at E5, in pigeon this accumulation was not detected before E8 (Figs. 1E, F; S1 G). In line with this result, the same ratios of surface areas occupied by mitochondria in the apical versus basal retinas were measured at E5 in chick and at E8 in pigeon (Fig. 1F). There was no bias toward apical localization of mitochondria in chick at E3, before the onset of neurogenesis (S1 H, I). The repartition of mitochondria identified with the MitoDsRed2 protein and by TEM is biased toward the apical retina at the beginning of cell differentiation, whereas mitochondria are more broadly distributed along the apico-basal axis in uncommitted progenitors.

**Figure 1:**
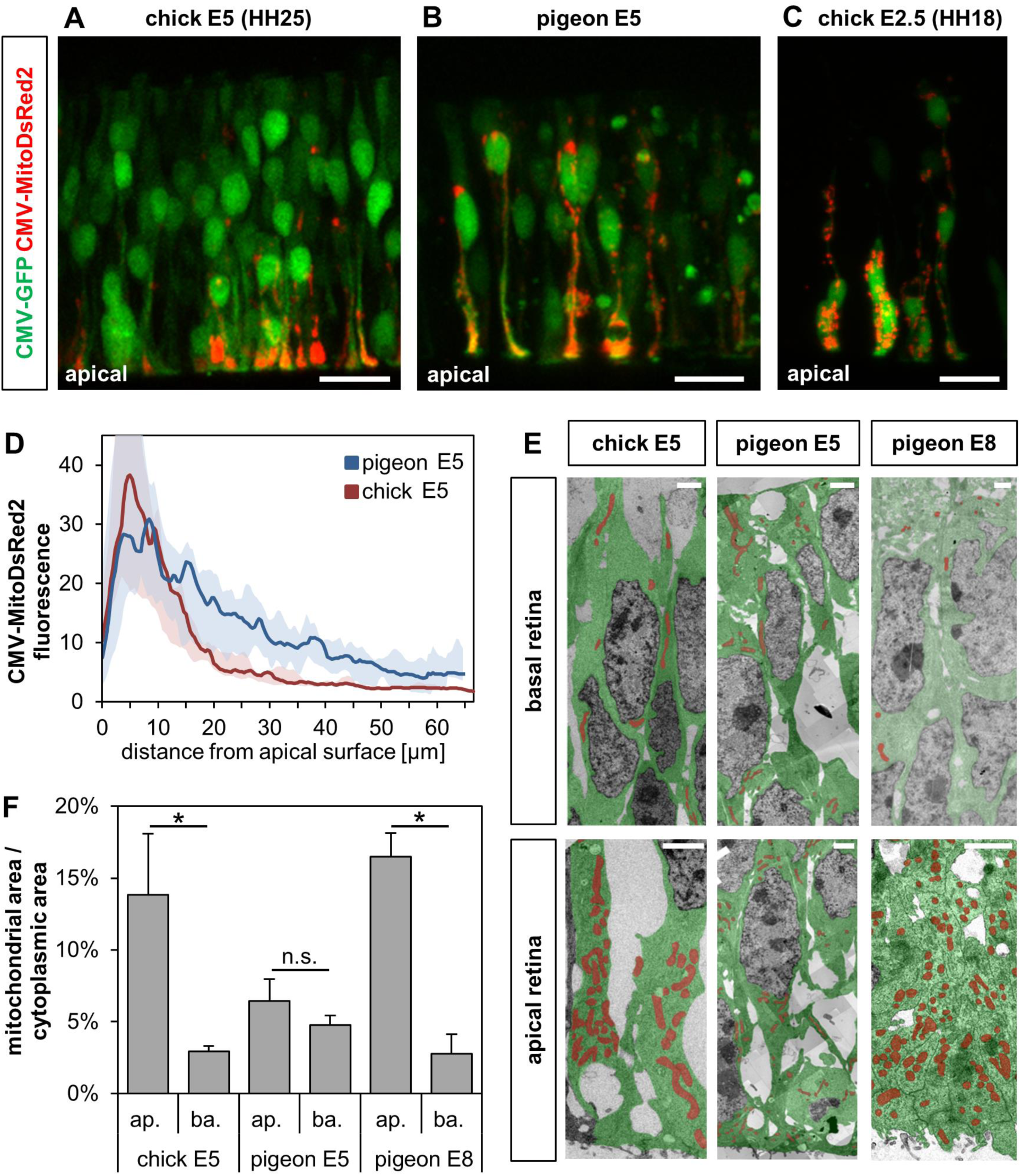
apical accumulation of mitochondria at the onset of cell differentiation. (A-C) Embryonic day 5 (E5) chick (A), E5 pigeon (B), E2.5 chick (C) retinas were co-electroporated with CMV-GFP and CMV-MitoDsRed2, and fluorescent cells were observed 24 h later by confocal microscopy. (D) Distribution of fluorescent mitochondria along the basoapical axis in chick and pigeon retinas at E5. The lines represent the mean values and the envelopes include the mean ± s.d. Mitochondria distribution was determined in 15 areas, ∼150 μm width, in 2 retinas of each species (E) E5 chick, and E5 and E8 pigeon retinas were processed for TEM. Images were stitched into a large mosaic to derive the apical and basal areas. Representative areas derived from these mosaics are shown, in which the mitochondria and the cytoplasm are artificially colored, respectively, in red and in green. (F) Ratio of the mitochondrial to cytoplasmic surface areas measured on the TEM picture mosaics. Histogram represents mean ± s.d., calculated from 2-4 mosaics from 2 retinas of each species, (*p<0.05; unpaired t-test; n=2-4 per group). Scale bars: 20 μm (A-C), 2-4 μm (E).

### Mitochondria in the RGC lineage

To determine when mitochondria started to accumulate apically, we monitored in real time the distribution of mitochondria in single Atoh7+ cells before their terminal mitosis (Movie S1). Apical accumulation of mitochondria roughly coincided with the upregulation of Atoh7 that occurs 8-15 h before the ultimate mitosis and marks the transition from pre-committed progenitors to cells committed to the RGC fate. *Hes5.3* is one of the earliest targets of the ATOH7 protein. The *Atoh7* and *Hes5.3* genes are co-expressed during an initial phase of selection of progenitors, when ATOH7 is expressed at a low level and progenitors are not yet committed to differentiation (Fig. 2A). HES5.3-mediated lengthening of the cell cycle is required for cells to enter the RGC lineage. Sibling cells of the ultimate mitosis express Atoh7 at high levels and extend axons indicating that both become RGCs (Chiodini et al., 2013). This view is supported by computational analysis of single-cell RNA sequencing data showing that cells expressing Atoh7 and cells expressing RGC markers (e.g., Nefm, Isl1, Pou4f3, Slit1, Stmn2, Chrnb3) belong to the same cluster (LB and JMM, unpublished data). The situation is different to that reported in zebrafish, where one Atoh7 progenitor generates one RGC and one non-RGC daughter cell (Poggi et al., 2005).

**Figure 2:**
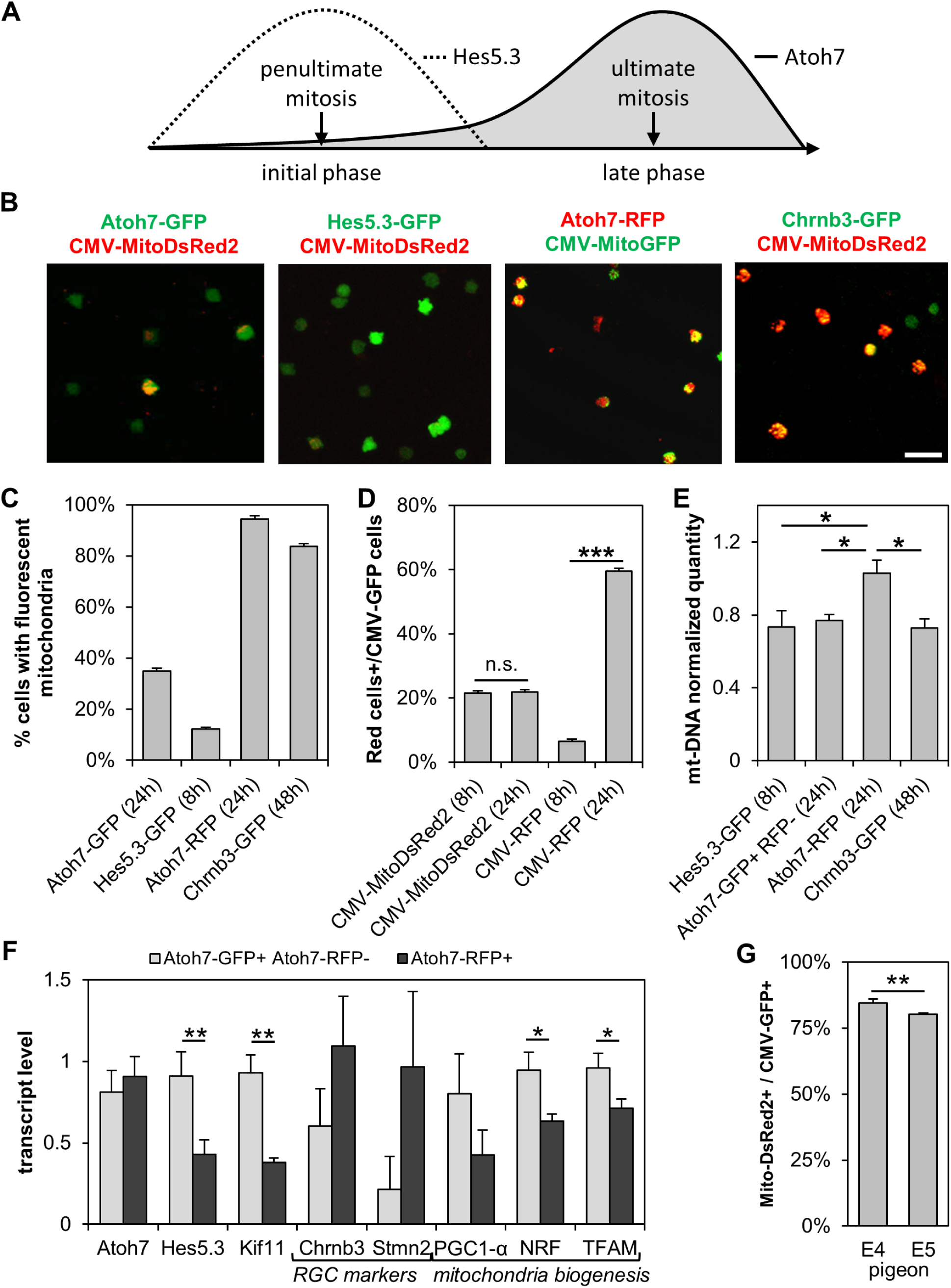
Mitochondria in the RGC lineage. (A) Schematic view of the time-course of Hes5.3 and Atoh7 expression during the penultimate and ultimate cell cycles. (B-F) E5 chick and (G) E4, E5 pigeon retinas were co-electroporated with reporter plasmids to identify different cell subsets. Retinas were disaggregated 8, 24 or 48 h later and subsets of cells were selected by FACS. (B-D) Cells were analyzed by confocal imaging. (C) Percentage of GFP+ or RFP+ cells with MitoDsRed2-or MitoGFP-labelled mitochondria. Data are presented as mean ± s.d. (Atoh7-GFP n=1879 cells, Hes5.3-GFP n=3531, Atoh7-RFP n=287, Chrnb3-GFP n=1571; all pairwise comparison show p<0.001; Chi-square test). (D) Percentage of GFP+ with labelled mitochondria and ratio of RFP+ to GFP+. Data are presented as mean ± s.d. (***p<0.001, n.s. not significant; Chi-square test; CMV-RFP 8h n=1017 cells, 24h n=1219, CMV-MitoDsRed2 8h n=3129, 24h n=2997). (E) The ratios of mt-DNA to g-DNA. Data are presented as mean ± s.d. (*p<0.05; unpaired t-test; n=3). (F) E5 retinas were co-electroporated with Atoh7-RFP and Atoh7-GFP. Cells were dissociated 24 h later. GFP+ RFP- and RFP+ cells were sorted by FACS. RNA was isolated and processed for RT-qPCR. Histogram shows mean ± s.d. (*p<0.05, **p<0.01; unpaired t-test; n=3). (G) Percentage of GFP+ cells with MitoDsRed2-labelled mitochondria. Data are presented as mean ± s.d. (**p<0.01, Chi square test; Pigeon E4 n=827, E5 n=4861). Scale bar: 20 μm

To monitor mitochondria accumulation along the pathway converting progenitors into newborn RGCs, E5 (HH25-26) retinas were electroporated with Atoh7-GFP, Hes5.3-GFP, Atoh7-RFP or Chrnb3-GFP in combination with a CMV-MitoDsRed2 or a CMV-MitoGFP reporter plasmid. Cells were dissociated 8, 24 or 48 h later and selected by fluorescence-activated cell sorting (FACS). Owing to developmental variations in the strength of the *Atoh7* promoter and to differences in the fluorescence properties between GFP and RFP (DsRed) (Strongin et al., 2007; Yanushevich et al., 2002; Yarbrough et al., 2001), the Atoh7-GFP plasmid identifies most of Atoh7 expressing cells, whereas Atoh7-RFP identifies only cells that express Atoh7 at high levels (See Fig. S4C in Chiodini et al., 2013). About 35% of cells identified with Atoh7-GFP and consisting of pre-committed progenitors and committed cells, contained MitoDsRed2-labelled mitochondria (Fig. 2B, C). The same proportion of Atoh7-GFP cells containing fluorescent mitochondria was found when *MitoDsRed2* was under the control of the Atoh7 promoter (Fig. S2), hence excluding inactivation of the CMV promoter in cells identified with Atoh7-GFP. The proportion of Atoh7+ cells containing MitoDsRed2-labelled mitochondria increased to ∼95% in committed cells identified with Atoh7-RFP (Fig. 2B, C). Consistent with the fact that committed cells express early RGC markers, ∼85% of cells identified with Chrnb3-GFP displayed mitochondrial labeling (Fig 2B, C). In contrast, only ∼12% of pre-committed progenitors identified with Hes5.3-GFP contained fluorescent mitochondria (Fig. 2B, C). The same proportion of CMV+ cells labelled with CMV-mitoDsRed2 was detected at 8h and 24h after electroporation (Fig. 2D) and we selected this short incubation period to diminish the likelihood of counting Hes5.3-GFP cells that already entered the RGC lineage. Like for Atoh7-GFP cells, the small fraction of cells identified with CMV-GFP and containing MitoDsRed2-labelled mitochondria (Fig. 2D) indicates that the low content of labelled mitochondria in Hes5.3+ cells (Fig. 2C) did not result from the lack of activity of the CMV promoter driving *MitoDsRed2*. These results raised the question of whether the CMV-GFP and Hes5.3-GFP reporters identify distinct cell populations. The pigeon retina helped to address this issue: at E4 and E5, cells are uncommitted and Hes5.3 is not expressed (Rodrigues et al., 2016), and yet ∼80% of cells identified with CMV-GFP contained fluorescent mitochondria (Fig. 2G), a characteristic of uncommitted cells (see below). Finally, if the MitoDsRed2 protein did not accumulate in mitochondria of Hes5.3+ and CMV+ cells, one could wonder why this fluorescent protein could hardly be detected in their cytoplasm. Chromophore maturation requires high DsRed2 concentration (Strongin et al., 2007; Yanushevich et al., 2002; Yarbrough et al., 2001). We surmise that the concentration threshold was reached for MitoDsRed2 proteins loaded into mitochondria, but not for proteins in the cytoplasm. In line with this idea, when cells were electroporated with CMV-RFP (DsRed2), the proportion of positive cells was much lower after 8h than 24h incubation, whereas the proportion of cells containing fluorescent mitochondria was the same (Fig. 2D).

To determine whether the ∼8-fold increase in the proportion of cells with fluorescent mitochondria between pre-committed progenitors and cells committed to the RGC fate was reflecting higher mitochondria content, we measured the ratio of mitochondrial DNA (mt-DNA) to genomic DNA (g-DNA). E5 retinas were co-electroporated with Atoh7-GFP and Atoh7-RFP plasmids, cells were dissociated 24 h later, and two subsets of cells were selected by FACS: one of GFP+ cells that did not express RFP and corresponding to Atoh7+ pre-committed progenitors, and the other one of RFP+ cells committed to the RGC fate. Likewise, pre-committed progenitors and newborn RGCs were selected using, respectively, Hes5.3-GFP and Chrnb3-GFP. Quantitative PCR (qPCR) analysis revealed higher ratios of mt-DNA to g-DNA in cells committed to the RGC fate than in pre-committed progenitors (Fig. 2E). A lower ratio of mt-DNA to g-DNA in RGCs identified with Chrnb3-GFP (Hernandez et al., 1995; Matter et al., 1995) than in committed cells identified with Atoh7-RFP probably reflects the fact that RGCs have axons filled with mitochondria (see Fig. 6C; Movie S2) that were lost during cell dissociation and sorting. The modest increase in the amounts of mt-DNA between pre-committed progenitors and committed cells did not match the strong increase in the proportion of cells containing fluorescent mitochondria (Fig. 2C). Nonetheless, we tested whether the augmentation of mt-DNA in committed cells identified with Atoh7-RFP reflected an increase of mitochondria biogenesis. E5 retinas were co-electroporated with Atoh7-GFP and Atoh7-RFP and cells were dissociated 24 h later. GFP+ RFP- and RFP+ cells were separated by FACS and processed for RT-qPCR (Fig. 2F). Hes5.3 and the mitotic marker Kif11 were expressed at lower levels in RFP+ cells, while RGC markers such as Chrnb3 and stathmin 2 (Stmn2) were upregulated. This molecular signature confirmed accurate discrimination between pre-committed progenitors and cells committed to the RGC fate. The upregulation of Atoh7 in committed cells was not detected in this assay, because at the time of RNA isolation, a fraction of RFP+ cells already turned off Atoh7 (Chiodini et al., 2013). Expression of PGC-1α, Nrf1 and Tfam, i.e., the main regulators of mitochondria biogenesis, were down-regulated in RFP+ cells (Fig. 2F) suggesting that cell cycle lengthening rather than mitochondria biogenesis led to the increase of mt-DNA content in committed cells (Fig. 2E).

### The disappearance of MitoDsRed2-labelled mitochondria in pre-committed progenitors results from a membrane potential decrease

The activation of *Hes5.3* by ATOH7 marks the onset of neurogenesis and the transition from uncommitted to pre-committed progenitors. Hes5.3 is turned on at ∼E4 in proliferating progenitors and turned off 8 to 15 hours before the ultimate mitosis (Fig. 3B inset; Chiodini et al., 2013). We monitored the proportion of Hes5.3+ cells containing fluorescent mitochondria at E4, E5 and E6 (Fig. 3A, B). The fraction of double-labelled cells abruptly decreased between E4 and E5 and reached a very low level at E6. In contrast, the proportion of uncommitted progenitors identified with Chrna7-GFP (Matter-Sadzinski et al., 1992) and containing MitoDsRed2-labelled mitochondria was maintained at a high level during the same period (Fig. 3C). We wondered whether the ∼10-fold decrease in the proportion of MitoDsRed2+ cells observed in the Hes5.3 subsets between E4 and E6 reflected a decrease in the number of mitochondria or a blockage in the import of MitoDsRed2 by mitochondria. To address this issue, we first did morphometric measurements on TEM images of Hes5.3+ cells selected by FACS at E4 and E6 (Figs. 3D; S3 A-E). Analysis at the single cell level did not reveal significant change in the total mitochondrial area per cell between E4 and E6 (Fig. 3D left panel). A modest increase in the number of mitochondria (Fig. 3D middle panel) and a decrease in the average area of individual mitochondria at E6 (Fig. 3D right panel) suggest that mitochondrial fragmentation could occur in Hes5.3+ cells. Likewise, we noted a slight decrease in the distance between mitochondria but no change in the circularity (Fig. S3 A, B). Overall, mitochondria density measured by TEM increased because of reduced cytoplasmic areas (Fig. S3 C, D). A similar increase in mitochondria density between E4 and E6 is detected with MTG (Fig. S3 F, G). Finally, similar amounts of mt-DNA were detected in Hes5.3+ cells at E4 and E5 (Fig. 3E). Next, we compared the expression of genes involved in mitochondria biogenesis and mitophagy in cells identified with Hes5.3-GFP at E4 and E5 (Fig. 3F). The accumulation of Hes5.3 and Atoh7 transcripts was significantly lower at E4 than at E5 confirming that our assay discriminated between *early* and *late* Hes5.3+ pre-committed progenitors. The master regulator of mitochondria biogenesis PGC1α was modestly down-regulated at E5, while NRF, TFAM, Ncor1 and the mitochondrial autophagy receptor Nix all remained unchanged. Taken together, our data suggest that the spectacular decrease in the abundance of MitoDsRed2-labelled mitochondria in Hes5.3+ cells was not accompanied by a decrease in the number of mitochondria. Then, we measured the mitochondrial membrane potential in retinal cells using fluorescent TMRM. TMRM labelling was lower in Hes5.3+ cells than in uncommitted progenitors identified with Chrna7-GFP (Fig. 4A), thus following the same trend as MitoDsRed2 (Fig. 3C). These results suggest a drop in mitochondrial imports of MitoDsRed2 in Hes5.3+ pre-committed progenitors resulting from a decrease in mitochondrial membrane potential.

**Figure 3:**
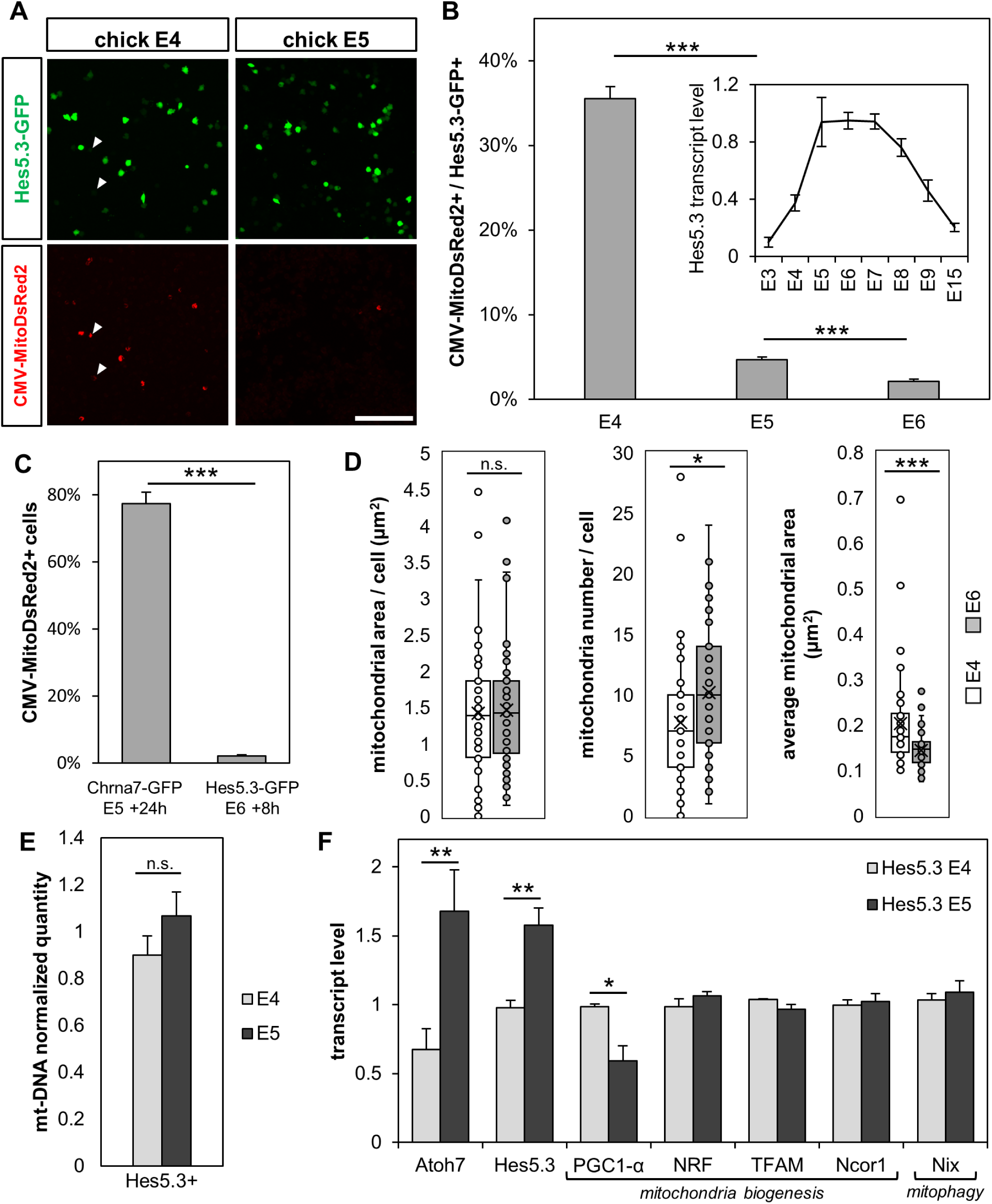
MitoDsRed2-labeling and mitochondrial count in Hes5.3+ pre-committed progenitors. (A, B) MitoDsRed2-labeling in Hes5.3+ cells. E4 (HH22-23), E5 (HH25-26) and E6 (HH28-29) retinas were co-electroporated with Hes5.3-GFP and CMV-MitoDsRed2. Retinas were disaggregated 8 h later and GFP+ cells were selected by FACS for confocal imaging (A). Arrowheads indicate MitoDsRed2 positive cells that are negative for Hes5.3-GFP (B) Percentage of GFP+ cells with mitochondria. Data are presented as mean ± s.d. (***p<0.001; Chi square test; E4 n=1126 cells, E5 n=3953, E6 n=3965). (B, inset) Accumulation of Hes5.3 transcripts measured by RT-qPCR as mean ± s.d. from biological triplicates. (C) Retinas were electroporated with Chrna7-GFP or Hes5.3-GFP and Mito-DsRed2-RFP to compare mitochondrial activity between uncommitted and pre-committed progenitors. GFP+ cells were sorted by FACS and the percentage ± s.d. of cells with mitochondria are shown (***p<0.001; Chi square test; Chrna7-GFP n=164 cells, Hes5.3-GFP n=3965). (D) Morphometric measurements of mitochondria in individual Hes5.3+ cells. E4 and E6 chick retinas were electroporated with Hes5.3-GFP. Dissociated GFP+ cells were sorted by FACS 8 h later and processed for TEM. Boxplot comprises median (central line) surrounded by 25^th^ and 75^th^ percentiles and mean indicated as cross. Whiskers show min and max values inside the 1.5 times interquartile range, and dots show all values including outliers (left p=0.7431, center p=0.0158, right p=0.0008; unpaired t-test; E4 n=51 cells, E6 n=55). (E) mt-DNA content relative to gDNA in Hes5.3+ cells. Cells were dissociated 8 h after electroporation, GFP+ cells were sorted by FACS and DNA was quantified. Data are presented as mean ± s.d. (p>0.05, unpaired t-test, n=3). (F) Transcript profiles in Hes5.3+ progenitors. Cells were dissociated 8 h after electroporation, GFP+ cells were sorted by FACS and RNA was quantified by RT-qPCR in triplicate. Data are presented as mean ± s.d., (*p<0.05, **p<0.01; unpaired t-test; n=3). Scale bar: 100 μm

**Figure 4:**
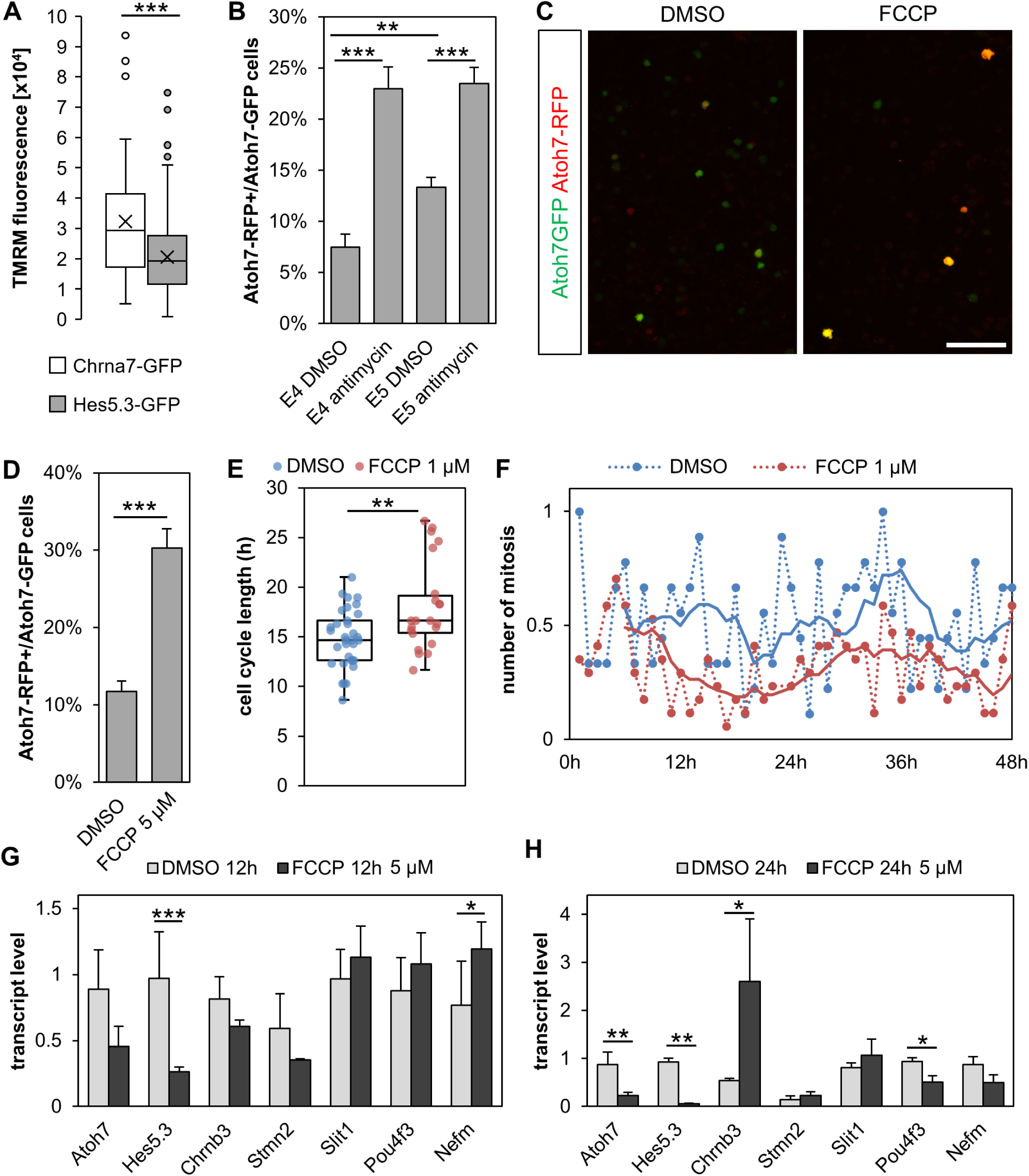
Low mitochondrial membrane potential promotes RGC genesis. (A) TMRM fluorescence in uncommitted vs. pre-committed progenitors. E4 or E5 retinas were electroporated with Chrna7-GFP or Hes5.3-GFP, respectively. Cells were dissociated 24 h (Chrna7-GFP) or 8 h later (Hes5.3-GFP) and stained with 1 nM TMRM. TMRM fluorescence was quantified in individual GFP+ cells as the sum of all pixel intensities (p<0.001; unpaired t-test; Hes5.3-GFP n=501 cells, Chrna7 n=51). (B-D) Antimycin A and FCCP increase the proportion of cells that enter the RGC lineage. (B) E4 and E5 retinas were electroporated with Atoh7-GFP and Atoh7-RFP and incubated with 36 μM Antimycin A or DMSO. Ratio of RFP+ to GFP+ cells are presented as mean ± s.d. (***p<0.001, **p<0.01, Chi square test, E4 DMSO n=427 cells, E4 Antimycin n=396, E5 DMSO n=1199, E5 Antimycin n=719). (C-D) E4.5 retinas were electroporated with Atoh7-GFP and Atoh7-RFP and incubated with 5 μM FCCP or DMSO. Cells were dissociated for confocal imaging (C) and cell counting (D). Ratio of RFP+ to GFP+ cells are presented as mean ± s.d. (***p<0.001, Chi square test, DMSO n=545 cells, FCCP n=635). (E, F) FCCP influences the cell cycle length. E4 retinas were electroporated with Chrna7-GFP and incubated with 1 μM FCCP or DMSO. (E) Cell cycle length from 22 cells (FCCP) and 33 cells (DMSO) tracked in real time from mitosis to mitosis. (F) Plot showing the average mitosis frequency as a function of time in 9 (DMSO) or 17 (FCCP) live imaging movies. Solid lines represent moving average over 6 periods. (G, H) FCCP influences RGC genesis. The right or left retinas from two embryos at E4 were incubated, respectively, with 5 μM FCCP or DMSO and processed for RT-qPCR analysis 12 h (G) or 24 h (H) later in triplicate. Scale bar: 50 μm

### Mitochondrial inhibitors affect cell cycle length and partially mimic the effect of HES5.3 on RGC fate commitment

HES5.3 increases the rate of conversion of pre-committed progenitors expressing Atoh7 at low levels into cells committed to the RGC fate and expressing Atoh7 at high levels (Chiodini et al., 2013). If this process involves a decrease of mitochondrial activity, one would expect that mitochondrial inhibitors should mimic, to some extent, the effect of HES5.3. To test this idea, E4 and E5 retinas were electroporated with Atoh7-GFP and Atoh7 RFP and incubated with Antimycin A, an inhibitor of the mitochondrial electron transport chain, or with FCCP. Compared to the controls, both inhibitors increased the ratio of RFP+ cells to GFP+ cells (Fig. 4B-D). In cell lines, FCCP or Antimycin A block G1-S transition and slow down progression of cells through the S phase (Han et al., 2008; Mitra et al., 2009; Schieke et al., 2008). Live imaging of retinal progenitors identified with Chrna7-GFP enabled us to test the effect of FCCP on the cell cycle progression. In the control retinas, the cell cycle lasted 14.9 h and cells divided during the 48 h of recording (Fig. 4E, F). Incubation of retinas with 1 μM FCCP induced the lengthening of the cell cycle (18.6 h) (Fig. 4E, F), while 5 μM FCCP blocked the cell cycle progression over a period of 24 h (Fig. S4A), an effect similar to that induced by the forced expression of Hes5.3 (see below, Fig. S6D). We reasoned that if the arrest of the cell cycle progression by FCCP led to an increase of Atoh7 promoter activity, it may also promote the production of RGCs. To test this idea, two E4 retinas from two separate embryos were incubated in culture medium containing FCCP while the two opposite retinas from the same embryos were incubated in medium with DMSO as a control. Retinas were processed for RT-qPCR analysis in triplicate 12 or 24 h later. A tendency towards increased expression of RGC markers (Slit1, Pou4F3, Nefm) after 12 h and the robust activation of Chrnb3 after 24 h in retinas incubated with 5 μM of FCCP (Fig 4G, H), but not with 1 μM (Fig 4B, C) suggests that reduced mitochondrial activity could promote RGC differentiation. However, the fact that Atoh7, Nefm and Pou4f3 were downregulated after 24 h of incubation (Fig. 4H) suggests that cells might fail to differentiate into mature RGCs. In the developing retina, Hes5.3 expression is downregulated and mitochondria recover their activity in cells committed to the RGC fate (see below, Fig. 6; Chiodini et al., 2013). In our experiment with FCCP, we did not reproduce a transient inhibition of mitochondrial activity similar to the one regulated by HES5.3, suggesting that the recovery of mitochondria activity in cells committed to the RGC fate is required to produce cells with a mature phenotype.

### HES5.3 promotes the decrease of mitochondrial activity

In order to evaluate the role of HES5.3 in the decrease of mitochondrial membrane potential, E5 chick retinas were electroporated with CMV-MitoDsRed2 and a β actin-Hes5.3:GFP expression vector or a CMV-GFP control vector. The forced expression of Hes5.3 led to a significant decrease in the proportion of cells with MitoDsRed2-labelled mitochondria 24 h later (Fig. 5A). Atoh7 is repressed by the Notch effector HES1 (Hairy1, ENSGALG00000002055) in early retina (Hernandez et al., 2007; Matter-Sadzinski et al., 2005) and gain- and loss-of-function experiments revealed the inhibitory effect of HES5.3 upon Hes1 (Fig. 5B, C). We asked whether the Hes1/Hes5.3 regulatory pathways could be involved in the decrease of mitochondrial activity. The forced expression of Hes1 in a subset of Hes5.3-expressing cells identified with Hes5.3-GFP at E5 resulted in an increase in the proportion of MitoDsRed2-positive Hes5.3+ cells 8 h later (Fig. 5D). By overcoming the inhibitory effect of Hes5.3 and repressing Atoh7 expression, HES1 kept cells in their uncommitted state (Matter-Sadzinski et al., 2005) and contributed to maintain MitoDsRed2-labelled mitochondria. We wondered how HES5.3 could trigger a decrease of mitochondrial activity in progenitors. Our Affymetrix analysis (see Materials and Methods and Chiodini et al. (2013)) revealed that HES5.3 could activate Cyp26A1, i.e., an enzyme controlling RA degradation, and repress a retinol dehydrogenase (RDH10), i.e., an enzyme involved in RA synthesis in Hes5.3+ pre-committed progenitors (Fig. 5E), suggesting that HES5.3 might be responsible for decreasing the level of RA. It is worthy of mentioning a coincidence between the disappearance of Cyp26A1 transcripts in the central area (HAA) of chick retina at HH28 (Da Silva and Cepko, 2017) and the turn-off of the *Hes5.3* promoter in a narrowly circumscribed central area at HH26-27 (Chiodini et al., 2013).

**Figure 5:**
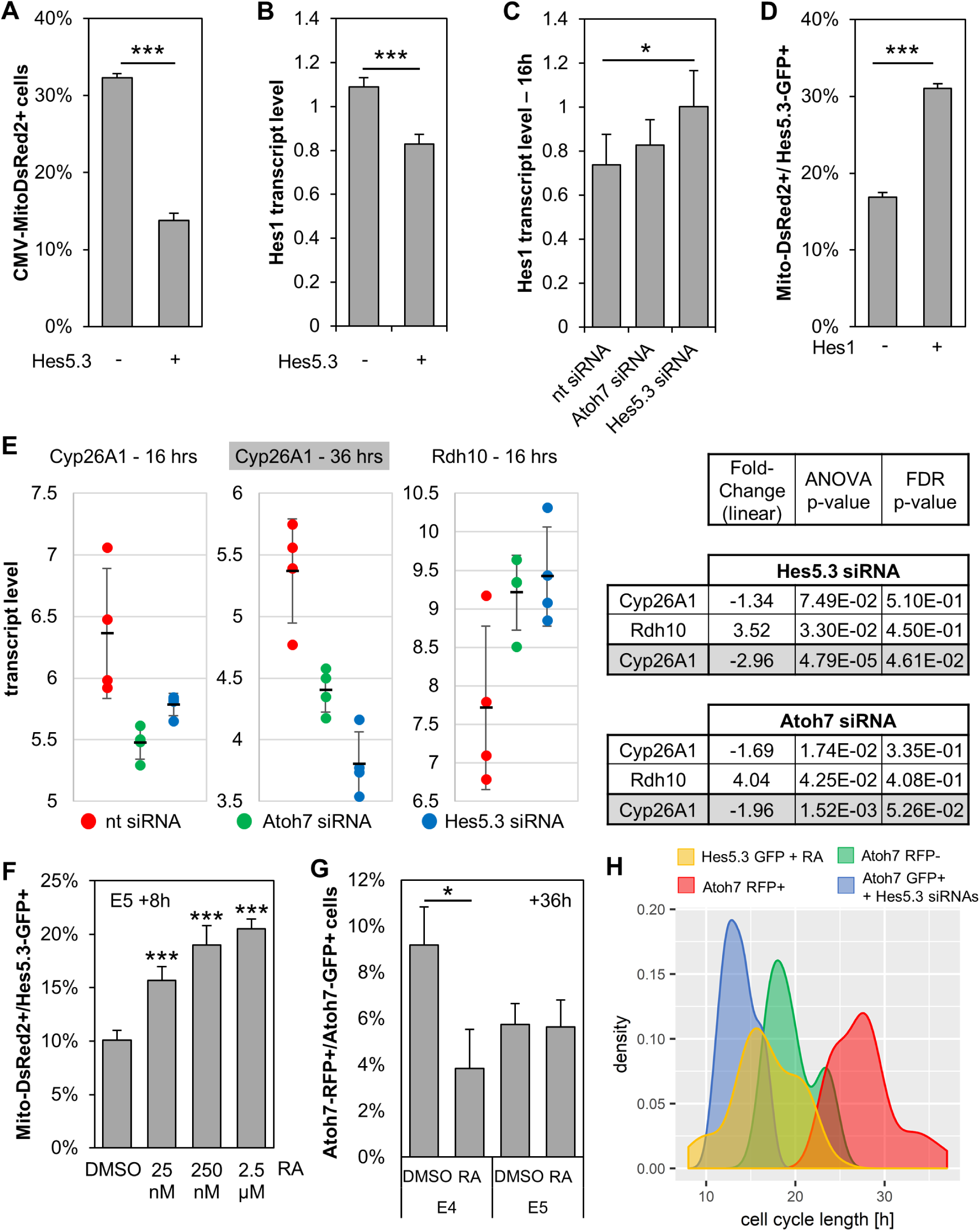
HES5.3 regulates mitochondrial activity via Hes1 repression and RA degradation. (A) Forced expression of Hes5.3 decreases the proportion of cells with MitoDsRed2-labelled mitochondria. E5 retinas were electroporated with β actin-Hes5.3:GFP or CMV-GFP, and CMV-MitoDsRed2. GFP+ cells were isolated by FACS 24h later. Percentage of GFP+ cells with mitochondria are presented as mean ± s.d. (***p<0.001; Chi square test; Hes5.3-GFP n= 96 cells, β actin-Hes5.3:GFP n=1421). (B, C) HES5.3 represses Hes1. (B) E3.5 retinas were electroporated with β actin-Hes5.3:GFP or CMV-GFP and processed for RT-qPCR analysis 24 h later. (C, E) E3.5 retinas were electroporated with siRNAs and Hes5.3-RFP. Retina fragments encompassing the edge of the expanding Hes5.3 domain were processed for RT-qPCR (C) or Affymetrix analysis (E) (16h, this paper; 36h, Chiodini et al., 2013). In B and C, data were obtained in four independent experiments and are presented as mean ± s.d. (***p<0.001, *p<0.05 Welch’s t test) (D) Forced expression of Hes1 increases the proportion of Hes5.3+ cells with MitoDsRed2-labelled mitochondria. E5 retinas were electroporated with Hes5.3-GFP alone or in combination with EMSV-Hes1. GFP+ cells were isolated by FACS 8 h later. Percentages of GFP+ cells with mitochondria are presented as mean ± s.d. (***p<0.001; Chi square test; ctrl n=4993 cells, EMSV-Hes1 n=5605). (E) Effects of Atoh7 and Hes5.3 siRNAs on the RA signaling pathway. For Rdh10, there is no effect at 36 h, probably due to HES5.3 stimulating the transcription of its own gene (Fior and Henrique, 2005; Chiodini et al., 2013). (F) RA increases the proportion of cells with MitoDsRed2-labelled mitochondria. E5 retinas were electroporated with Hes5.3-GFP and CMV-MitoDsRed2 and incubated with RA or DMSO. Cells were dissociated 8 h later. Percentages of GFP+ cells with mitochondria are presented as mean ± s.d. (***p<0.001, Chi square test, DMSO n=1129 cells, 25 nM RA n=772, 250 nM RA n=484, 2.5 μM RA n= 9 0) (G) RA decreases the proportion of cells that enter the RGC lineage. Retinas at E4 and E5 were electroporated with Atoh7-GFP and Atoh7-RFP and incubated with 2.5 μM RA or DMSO. Cells were dissociated 36 h later. Ratio of RFP+ to GFP+ cells in the presence or absence of RA. Data are presented as mean ± s.d. (*p<0.05; Chi-square test; E4 DMSO n=305 cells, E4 RA n=130, E5 DMSO n=678, E5 RA n=384). (H) Four E4 chick retinas were electroporated with Hes5.3-GFP, 2.5 μM RA as added and retinas were processed for Time-Lapse imaging 24h later. 64 cells were tracked from mitosis to mitosis. Density plot of cell cycle length (data in Fig. S6B) showing that cell cycle of a subpopulation of Hes5.3-GFP cells treated with 2.5 μM RA (yellow) is similar to that of Atoh7-GFP cells treated with Hes5.3 siRNA (blue) and shorter than that of cells expressing Atoh7 at low (green) or high levels (red). Cells expressing Atoh7 at low level (green) and cells expressing Hes5.3 constitute the same subset (Chiodini et al., 2013).

### RA favors mitochondrial activity over glycolysis

The negative effect of HES5.3 on RA is interesting in light of a recent study reporting the inhibitory effect of RA on RGC genesis (Da Silva and Cepko, 2017). We wondered whether this was related to the decrease of mitochondrial activity. Indeed, incubation of E5 retinas with RA increased the proportion of Hes5.3+ cells containing MitoDsRed2-labelled mitochondria in a dose dependent manner (Fig. 5F). Moreover, when E4 retinas were co-electroporated with Atoh7-RFP and Atoh7-GFP and incubated with 2.5 µM RA for 36 h, the ratio of RFP+ to GFP+ cells was lower than in the control (Fig. 5G). It appears that RA might increase mitochondrial activity and diminish the chance for progenitors to enter the RGC lineage. In order to assess the role of RA, we compared the RNA-Seq transcriptomes of Hes5.3+ cells isolated from E5 retinas incubated either with 2.5 µM RA or with DMSO for 8 h (Fig. S5). RA treatment reduced expression of genes encoding glycolytic enzymes, and more particularly, the lactate dehydrogenase A (LDHA) which plays a key role in aerobic glycolysis (the Warburg effect) (Liberti and Locasale, 2016). RA also reduced expression of *Rcan1* which has been linked to mitophagy and glycolytic switch (Ermak et al., 2012). On the other hand, RA increased expression of genes which could positively influence mitochondrial activity. For example, the ephrins B2 (EFNB2) and its receptor (EPHB2) can activate the mitochondrial translocation of Sirt3 (Jung et al., 2017), i.e., a deacetylase that activates mitochondrial function (Sun et al., 2018). Taken together, our data suggest that RA favors mitochondrial activity at the expense of aerobic glycolysis and reduces the chance that Atoh7+ / Hes5.3+ pre-committed progenitors convert into cells committed to the RGC fate.

### RA accelerates the cell cycle progression

Having shown that the role HES5.3 played in RGC fate commitment could involve RA degradation, we sought to identify the commitment point at which decreased mitochondrial activity was required. Hes5.3 inhibition accelerates the cell cycle (Chiodini et al., 2013) and we wondered whether incubation of retina with RA could have a similar effect. E4 retina explants were electroporated with Hes5.3-GFP, incorporated in a 3D matrix of collagen and incubated with 2.5 µM RA for 24 h. We monitored progression of fluorescent cells through the cell cycle by time-lapse imaging (Figs. 5H; S6A, B). The cell cycle length of 64 Hes5.3+ progenitors that we tracked from mitosis to mitosis lasted 8 to 24 hr. They divided in two subsets (Fig. 5H): one displaying cell cycle length within the range of cells identified with Atoh7-GFP, and the other having shorter cycle comparable to that of cells in which Hes5.3 expression was inhibited by siRNAs. We interpret this as evidence that RA accelerated the cell cycle shortly after the onset of Hes5.3 expression but had no effect at later stages. This is consistent with the fact that RA decreased the ratio of Atoh7-RFP+ to Atoh7-GFP+ cells at E4 but not at E5 (Fig. 5G) and that RA decreases RGC genesis in the central area (HAA) but not in peripheral zones that develop at later stages (Da Silva and Cepko, 2017). If increased mitochondrial activity could accelerate the cell cycle, we wondered what the effect of a decrease of mitochondrial activity resulting from the forced expression of Hes5.3 would be (Fig. 5A). Time lapse imaging of fluorescent cells in E4 retinas electroporated with a β-actin-Hes5.3:GFP expression vector revealed that cell cycles lasted ≥ 30 hrs because of much delayed apex-directed migration prior to mitosis on the apical surface (Fig. S6D). Taken together, these results suggest that after a lapse of low mitochondrial activity in Hes5.3+ pre-committed progenitors, the downregulation of Hes5.3 is required for the apical accumulation of active mitochondria during the period preceding the ultimate mitosis.

### Mitochondria relocate in newborn RGCs

After their ultimate mitosis, newborn RGCs are paired along the apical surface where they reside for ∼15 h. This arrest coincides with the robust accumulation of the ATOH7 protein (Fig. 2A; Chiodini et al., 2013). To monitor in real-time mitochondria distribution in RGCs, E5 chick retinas were electroporated with Atoh7-GFP or the early RGC marker Chrnb3-GFP and Atoh7-MitoDsRed2 or CMV-MitoDsRed2 reporter plasmids. Cells imaged on the apical side were initially bipolar with a short apical process (Fig. 6A cell 1), and shortly after the terminal mitosis an axon started to grow at the basal pole. When the growth cones reached the basal surface, they turned at right angle (white arrowheads in Fig. 6A cell 2, and Fig. 6C) and navigated along the basal surface toward the head of the optic nerve. At this time, RGC somas slowly and steadily migrated toward the basal surface where they established the ganglion cell layer (Fig. 6A cells 2 and 3, B, C; Chiodini et al., 2013). Mitochondria were restricted to the apical process in newborn RGCs, and this localization persisted until RGC soma migrated to the basal side. More precisely, mitochondria accumulated at the endfoot of the apical process (Fig. 6A cells, 1’, 1’ 1”, Fig 6C), with rare events of transient distribution across the entire apical process (not shown). During migration of the soma to the basal surface, mitochondria were redistributed around the nucleus (Fig. 6A cell 2, Fig.6B). First, they clustered apically in the soma with transient relocation to the basal side (Fig. 6B yellow arrowheads). As the cell progressed toward its final position, mitochondria were more often present either at both poles (Fig. 6A cell 2, C 8h red arrowheads) or on the basal side of the soma only (Fig. 6B 8h20 blue arrowhead). Finally, mitochondria moved into the axon with a lag of a few hours after the axon had started to grow (Fig. 6C 8h vs. 12h yellow arrowheads). In sum, there is a strong apical accumulation of mitochondria in newborn RGCs, followed by the relocation of mitochondria from the apical to the basal poles while RGCs move to the basal side and extend their axons on the basal surface (Movie S2).

**Figure 6:**
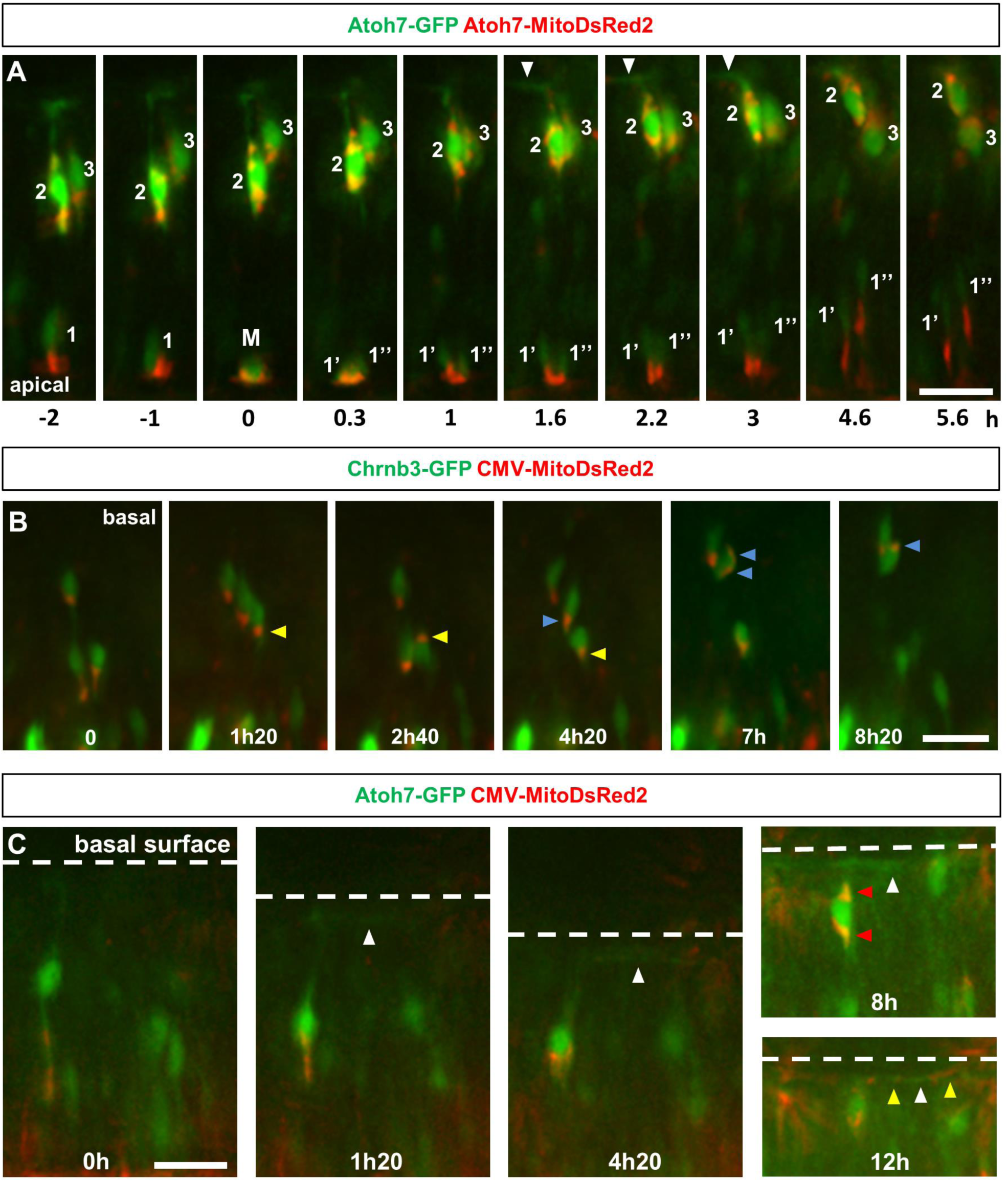
mitochondria relocate in newborn RGCs. (A-C) Stills from time-lapse movies in the green and red channels. Electroporated E5 chick retinas were incorporated in a collagen matrix and imaged 24 h later (t_0_). (A) Cell 1: the stills show mitochondria accumulation at the apex of an Atoh7+ cell before the last mitosis (M). Mitochondria are equally distributed between the two daughter cells (1’ and 1”) Cells 2 and are newborn RGCs reaching the basal side (the white arrowheads show an axon). The relative differences in fluorescence intensities between the cell 1 and the cells 2 & 3 are consistent with the fact that, while activity of the Atoh7 promoter increased before the ultimate mitosis, it reached peak level well after the last cell division. Cells strongly labeled with the ATOH7 antibodies are unlabeled with BrdU (Fig. S3, Chiodini et al., 2013). (B, C) Mitochondria localization in newborn RGCs expressing Chrnb3 (B) or Atoh7 (C). (B) Mitochondria translocate from the apical to the basal side (blue arrowheads) or undergo back and forth movements (yellow arrowheads) during the basal migration of the soma. (C) The white arrowheads show an axon filled with mitochondria (yellow arrowheads at 12 h). (C, 8 h) A RGC reaching the basal aspect with mitochondria at both poles of the cell body (red arrowheads). Scale bars: 20 μm (A, B, C).

## DISCUSSION

The number of mitochondria per cell likely reflects the fine balance between different processes including biogenesis, mitophagy as well as cytokinesis itself. Both asymmetric segregation of mitochondria in daughter cells and mitophagy could unbalance mitochondria counts. In mouse retina, mitophagy creates this unbalance in mitochondrial mass between neuroblasts and differentiating RGCs (Esteban-Martinez et al., 2017). In chick, no mitophagy was detected in the RGC lineage and mitochondria apportioning in daughter RGCs was roughly equal. In chick and pigeon early embryonic retinas, mitochondria biogenesis kept pace relatively well with the rapid expansion of the pool of retinal progenitors. The ∼4-fold decrease in the ratio of mt-DNA to g-DNA between E4 and E8 remained modest compared with the ∼100-fold increase in cell number during this period (Cherix et al., submitted). The decrease of mitochondrial mass in the course of retina development correlates with a marked decrease of mitochondrial activity except in RGCs. The lengthening of the cell cycle of Hes5.3+ pre-committed progenitors mitigates the decline of mitochondrial content, while the number of mitochondria per cell increases in newborn RGCs despite a decrease in the expression of genes involved in mitochondria biogenesis. A drop in mitochondrial membrane potential is the unique evidence of a metabolic shift in Hes5.3+ pre-committed progenitors. This parsimonious model of regulation should aim at preserving mitochondria to assure that they regain activity shortly before the ultimate mitosis and in newborn RGCs. How a disruption of the proton gradient across the inner mitochondrial membrane could be associated to cell commitment? It appears that the property of ATOH7 to decrease RA level through the activation of Hes5.3 and Cyp26A1 and, thereby, favoring glycolysis over OXPHOS is instrumental in establishing the link between cell commitment and metabolism (Fig. 7). This link helps to gain understanding of why high levels of RA or the inhibition of Hes5.3 expression can reduce the production of RGCs in the retina (Chiodini et al., 2013; Da Silva and Cepko, 2017). In pigeon, robust and sustained mitochondrial activity in early retina suggests that RA level could remain high over a longer period of time than in chick, thereby contributing to postpone RGC determination and differentiation until the end of the period of cell proliferation.

**Figure 7:**
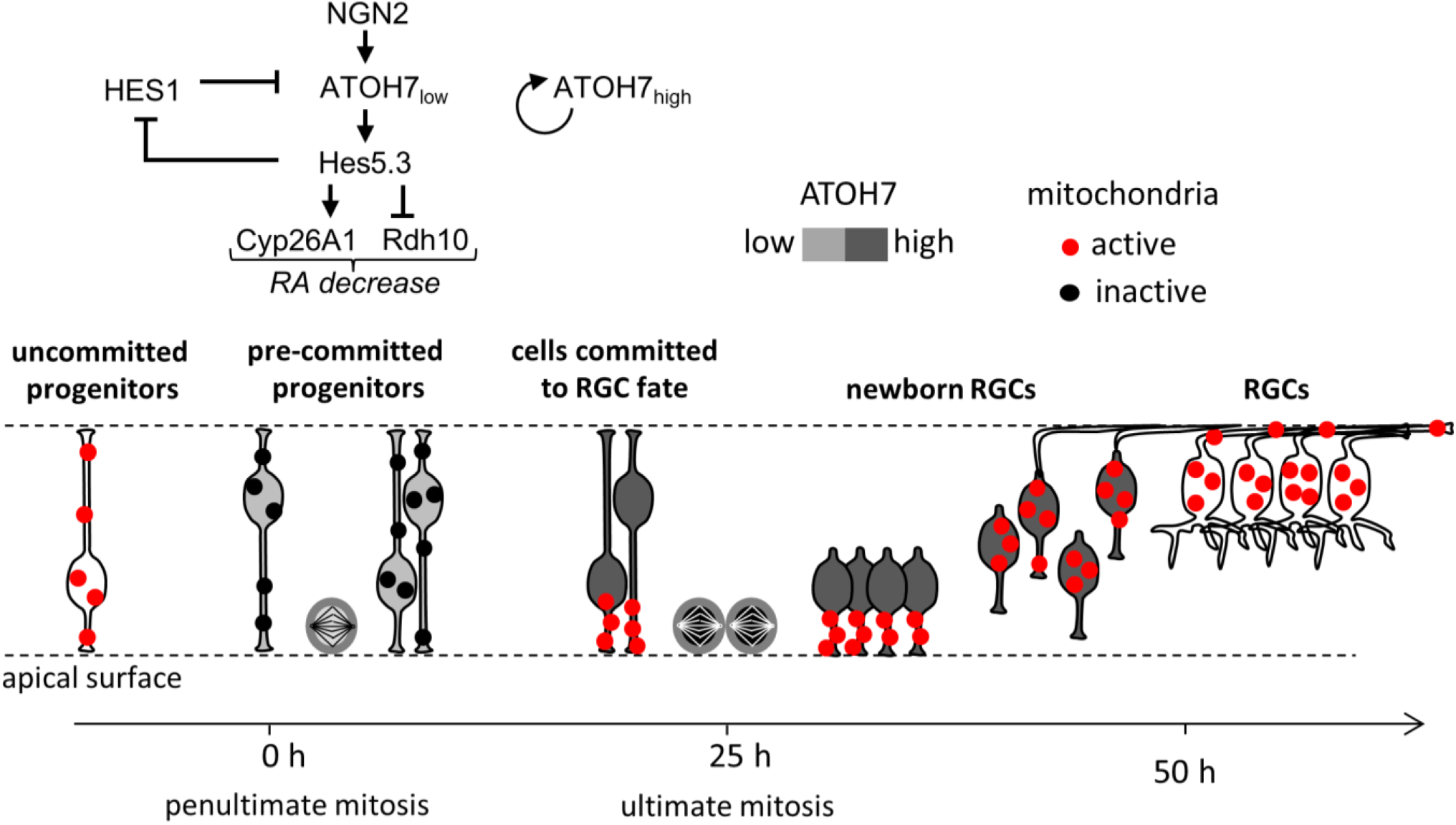
Model describing associations between key transcription factors, the RA signaling pathway and changes in mitochondrial activity. ATOH7 activates Hes5.3, which, in turn, induces a decrease in mitochondrial activity via the RA signaling pathway. This metabolic shift slows down the cell cycle progression and is a necessary step for ATOH7 to induce RGCs.

We wondered how a transient decrease of mitochondrial activity can influence the RGC fate determination, considering that the upregulation of Atoh7 and the lengthening of the cell cycle are required to achieve this goal. RGCs are the first neurons born in the retina and they are specified at stages when the vast majority of retinal progenitors divide at a high rate with cell cycle lasting less than 15 h. Uncommitted progenitors have a high content of active mitochondria and their low lactate production suggest that they predominantly utilize OXPHOS for energy supply (Cherix et al., submitted). Rapid rate of mitosis justifies high energetic requirement (Carbognin et al., 2016; Miettinen and Bjorklund, 2017), even though progenitors need a supply of glycolytic intermediates essential for anabolic reactions during cell division. The transient decrease in the number of active mitochondria into RGC-biased pre-committed progenitors probably does not result from a downturn in oxygen and substrate supply, because a large subset of uncommitted progenitors at the same location maintain a high content of active mitochondria. Our data suggest that the HES5.3-mediated inactivation of mitochondria can be a strategy for slowing down cell cycle progression in response to a decrease in ATP synthesis. Although we do not know yet whether there is a decrease of ATP level in Hes5.3-expressing cells, previous studies (Mitra et al., 2009) show that mitochondrial uncoupling with FCCP slows down the cell cycle progression. In the retina, we show that FCCP treatment lengthens the cell cycle progression and promotes RGC genesis. Taken together, our data suggest that cell cycle lengthening is a commitment point that requires a transient metabolic shift towards aerobic glycolysis.

It appears that aerobic glycolysis is required for RGC production in chicken and mouse. However, each species follows a distinct regulatory pathway to achieve this purpose. The fact that newly generated mouse RGCs contain fewer mitochondria than retinal neuroblasts and that low mitochondrial content is required for normal axon growth is one out of several differences in the production of RGCs between birds and rodents (Esteban-Martinez et al., 2017). In chick, mitochondria adopt very specific distribution in newborn RGCs, with their apical accumulation before the onset of axogenesis and their migration towards the basal side when the axon starts to grow. What could be the meaning of the apical accumulation of mitochondria in committed RGCs shortly before their ultimate mitosis? Mitochondria regained activity in committed cells and their apical position could facilitate ATP supply for bringing back cells in G2 on the apical surface and for mitotic spindle formation and myosin ring contraction to complete cytokinesis. Oxygen supply is another reason that may justify mitochondria accumulation on the apical surface. When RGCs differentiate, the pecten oculi is not yet developed and oxygen comes from the choroid, making an apical oxygen gradient plausible. This could explain why newborn RGCs with high levels of ATOH7 and with growing axons pause for ∼15 h on the apical surface after their ultimate mitosis (Fig. 7; Chiodini et al., 2013).

Significant differences between RGC production in chick and mouse could arise from variations in the regulation of *Atoh7* (Skowronska-Krawczyk et al., 2009) and from the possible absence of an ortholog of *Hes5.3* in mammals (Fior and Henrique, 2005). As a consequence, while in chick, ∼80% of Atoh7+ pre-committed progenitors develop into RGCs, the proportion drops to ∼15% in mouse where the majority of Atoh7-expressing cells become photoreceptors (Brzezinski et al., 2012; Skowronska-Krawczyk et al., 2009; Yang et al., 2003). It would be interesting to know whether the metabolic shift toward aerobic glycolysis implemented in birds for optimizing the production of RGCs, serve a similar function for cell type specification and the generation of cell type diversity within other areas of the central nervous system.

## MATERIALS AND METHODS

### Animals

Chick embryos from a White Leghorn strain (UNIGE Animal Resources Centre) were staged according to Hamburger and Hamilton (1951). Fertilized pigeon eggs were supplied by Philippe Delaunay (Pigeonneau de la Suisse Normande, Croisilles, France). Experimental procedures were carried out in accordance with Federal Swiss Veterinary Regulations. Eyes were dissected in DPBS (ThermoFisher) and the surrounding retinal pigment epithelium (RPE) was removed few minutes before electroporation.

### Reporter and expression plasmids

The Chrnb3 (β3 nAChR)-GFP plasmid derives from a β3 nAChR-CAT00 plasmid (Hernandez et al., 1995) The restriction fragment beginning 128 bases (− 128, phI) 5′ of the β initiator ATG and extending −271 (EcoRI, fragment RS, 143 bp in length) as inserted into the smal site of pCAT00, immediately upstream of the CAT gene. pCAT00 (Crossley and Brownlee, 1990) contains two synthetic polyadenylation sites upstream of the SmaI site and has low-background activity in retinal cells (Hernandez et al., 1995). The Atoh7(Ath5)-GFP, Hes5.3-GFP, Chrna7-GFP plasmids and the Atoh7-RFP (DsRed2) plasmid derive, respectively, from pEGFP-C1 and pDsRed2-N1 plasmid (Clontech). The fragment bounded by the AseI and NheI restriction sites was deleted from pEGFP-C1 and pDsRed2-N1 and the upstream sequences of Atoh7 (2220 bp in length, Hernandez et al., 2007), Hes5.3 (1960 bp in length, Chiodini et al., 2013) or Chrna7 (406 bp in length, Matter-Sadzinski et al., 1992) were subcloned in the proper orientation at appropriate sites in the vectors. The pEMSV plasmid was used to express the *cAtoh7, cHES1* (Hairy1, ENSGALG00000002055) in co-transfection and electroporation experiments (Hernandez et al., 2007; Matter-Sadzinski et al., 2005). The β actin-Hes5.3:GFP expression vector (Fior and Henrique, 2005) is a gift from D. Henrique. The CMV-MitoDsRed2 (pDsRed2-Mito) and CMV-MitoGFP (pAcGFP1-Mito) plasmids are from Clontech. Atoh7-MitoDsRed2 was designed using UGene software (Unipro). The GFP sequence was excised from Atoh7-GFP plasmid using AgeI and KpnI restriction enzymes. The MitoDsRed2 sequence from the CMV-MitoDsRed2 plasmid was amplified with PCR and extremities were digested with XmaI and KpnI restriction enzymes. Ligation was performed using Quick Ligation Kit.

### MitoTracker

Retinas or dissociated cells were incubated with MitoTracker Green FM (MTG, ThermoFisher) at 150 nM concentration in DMEM (Amimed) for 30-40 minutes in a 37°C incubator with 5% CO_2_. After washing with DMEM, cells were imaged directly under confocal microscope at a constant 37°C temperature.

### Retina electroporation

Stripped eyes were electroporated in electroporation cuvettes (BT 640, BTX) with the reporter plasmids CMV-GFP, Atoh7-GFP, Atoh7-RFP, Hes5.3-GFP or Chrnb3-GFP at 0.5 µg/µl, CMV-MitoDsRed2 or CMV-MitoGFP at 0.1 µg/µl, and Atoh7-MitoDsRed2 at 2 µg/µl. Expression vectors EMSV-Atoh7, EMSV-Ngn2, EMSV-es1, and β actin-Hes5.3:GFP were used at 0.5 µg/µl. Electroporation was performed in 100 µl using 5 pulses of 5 V and 50 ms, separated by 1 second interval with BTX ECM830 electroporator.

### Tissue culture

Electroporated retinas were cultured in DMEM (ThermoFisher) complemented with 10% Fetal Bovine Serum (ThermoFisher) and 1% Penicillin-Streptomycin (ThermoFisher) for 8 h, 24 h, or 48 h at 37°C in an incubator with 5% CO_2_.

### Tissue dissociation

After 8 h or 24 h of tissue culture, electroporated retinas were washed with HBSS lacking Ca^2+^ and Mg^2+^ (ThermoFisher) and dissociated using Trypsin 0.05 % (ThermoFisher) for 15-20 minutes at 37°C. Reaction was blocked by addition of 10% Fetal Bovine Serum (ThermoFisher).

### Fluorescence Activated Cells Sorting

After dissociation, cells were pelleted and resuspended in DMEM (Amimed). Cells were sorted by FACS Aria II for GFP positive cells, or with FACS Astrios for GFP and RFP positive cells. Cells were pelleted and resuspended at 1-2 x 10^6^ cells/ml. 100-200 µl cell suspension were plated on a poly-dl-ornithine (Sigma-Aldrich) coated permanox chamber slide (Lab-Tek). Cells were left 30 minutes at 37°C and 5% CO_2_ for adhesion. Cells were fixed with 4% paraformaldehyde for 20 minutes. Finally, after DPBS washing, DABCO (Sigma-Aldrich) was added and the slide was sealed with coverslip for imaging.

### RT-qPCR and mt-DNA quantification by qPCR

RNA from FACS sorted cells or from whole retinas was extracted using TRIzol reagent (ThermoFisher) according to product manual in triplicate. Primers were designed using NCBI primer blast and Primer3 websites. Primers were ordered from Microsynth. Primers were tested using RNA from total retina extracted with TRIzol. RNA quantification was done with spectrophotometer and Qubit 2.0 (ThermoFisher) and RNA quality was checked using BioAnalyzer 2100 (Agilent). RT was performed using Takara PrimeScript reverse transcriptase prior to qPCR. In samples with delta-Ct values > 0.5 across the three technical replicates, the most extreme value was removed. Quantities were calculated as 2^(Ctmin-CT). The Average-Quantity was calculated for each sample. geNorm (Vandesompele et al., 2002) was used to determine the best normalization genes. A normalization factor was calculated as Normalization-Factor=Geometric Mean [Average-Quantity of Normalization Genes] and was used to calculate the normalized quantity: Normalized-Quantity=Average-Quantity/Normalization-Factor. The fold change was calculated as: Fold-Change=Mean[Experiment condition] / Mean[Baseline condition]. P-values were calculated using a Welch t-test to compare baseline and experiment samples (in log2 scale). DNA of FACS sorted cells or dissected retinas was extracted according to DNeasy Blood & Tissue Kit (Qiagen). Primers were designed for targets on g-DNA and mt-DNA. qPCR was performed and g-DNA used for normalization. RT-qPCR and qPCR were performed using the primers listed in Table S1. All specific primers for chick and pigeon listed in Table S1 were tested for efficiency.

### Confocal imaging

Intact retinas were fixed with 4% paraformaldehyde for 20 minutes and were mounted on concave glass slides. Dissociated or FACS sorted cells were plated on permanox chamber slides (Lab-Tek) coated with poly-dl-ornithine, left 30 minutes for adhesion in a 37°C 5% CO_2_ incubator, and then fixed 20 minutes with 4% paraformaldehyde. Coverslips were sealed after addition of DABCO with 50% glycerol (Sigma-Aldrich) in DPBS (ThermoFisher). Imaging was done with a Leica Sp5 Laser scanning Confocal microscope in photon counting mode using a Leica 20x multi-immersion objective (0.7 N.A.) in Leica type F-type immersion oil of refractive index 1.518. An Argon laser was used for 488 nm GFP excitation and DPSS561 for 561 nm RFP excitation. Optical sections of 1 μm were acquired with a 2-4x line accumulation.

### Time-Lapse imaging

To monitor interkinetic nuclear migration (INM) and mitosis at the retina equator, electroporated retinas were placed on a 35-mm glass bottom dish (Pelco, Wilco Wells) and incorporated in collagen prepared as follow: 100 mg of rat tail collagen type VII (Sigma-Aldrich) was dissolved in 6ml of 16.7 mM acetic acid overnight at 4°C. 18 ml of double distilled water was added and the solution was dialyzed overnight against 4.2 mM acetic acid at 4°C. Finally, 550 µl of this reconstituted collage, 80 µl of 4.2 mM acetic acid, 100 µl of DMEM 10x (Amimed) and 100 µl of 0.28 M sodium bicarbonate were mixed. The dish was incubated at 37°C to allow collagen polymerization. Culture medium composed of DMEM without phenol red complemented with 10% fetal bovine serum and 1% Penicillin-Streptomycin was added on top of the collagen matrix. In experiments using TMRM (Image-iT TMRM Reagent, ThermoFisher), FCCP (Sigma-Aldrich) or retinoic acid, drug was added directly to culture medium and reached retinas through collagen in 15-20 minutes. Retinas were imaged 24 h after electroporation using a Leica Widefield AF6000LX microscope with a Leica 40x dry long-distance objective (0.55 N.A.) for up to 72 h in the Green and Red channel using Leica GFP and Rhodamine filter cubes. Stacks of images separated by 1 μm for a total of 40-60 microns were used with a time interval of 20 minutes, 2 minutes or 20 seconds. Data were saved as *.lif file.

### Transmission Electron Microscopy (TEM)

Retinas or pellets from FACS sorted cells were fixed >24 h in 3.7% formaldehyde, 1% glutaraldehyde 0.1 M Sodium phosphate monobasic, 0.07 M NaOH in H_2_O at 4°C. Tissue were washed with 0.1 M ice cold sodium cacodylate before incubation 10 minutes in 0.8% K_3_Fe(CN)_6_ in 0.1 M sodium cacodylate. The tissue was fixed in 1% osmiumtetroxide, 0.8% K_3_Fe(CN)_6_ in 0.1 M sodium cacodylate pH 7.4 for 90 minutes. The tissue was washed with sodium cacodylate and water, and stained 2h with 1% uranyl acetate in dark. Tissue was dehydrated in ethanol baths and incubated in propylene oxide 100% for 45 minutes, followed by propylene oxide/epon 1:1 incubation overnight. Tissue was embedded in epoxy resin consisting of 48% Agar 100, 18% DDSA, 31% MNA and 3% BDMA. 80-90 nm sections were produced using a Leica UCT ultra-microtome, and were recovered on TEM grids. Staining was done by incubation 5 minutes in Acetone / Uranyl Acetate 1:1 solution followed by 10 minutes in lead citrate solution. TEM images were acquired using a Tecnai G2 Transmission Electron Microscope. Mitochondria and cytoplasm were delimited in Photoshop CS2 (Adobe), and areas were measured in imageJ/Fiji. The area ratio of mitochondria to cytoplasm was calculated.

### Image processing and morphometry

3D blind deconvolution of Time-Lapse dataset was done with Autoquant X3 software (MediaCybernetics). Deconvolution was used to reduce out of focus noise and increase the contrast. Further image processing was done using ImageJ and Fiji softwares. Cell cycle length was determined by following cells from mitosis to mitosis in 4D live imaging movies. Quantification of mitosis frequency (as displayed in figure 4F and S4 A) was done by marking mitosis in every frame of the 4D movie. Each movie represented a total volume of 5.15E6 μm^3^ (293 μm x 293 μm x 60 μm) that included the optical cross section of the retina equator. The average number of mitosis for each time frame was calculated across all movies for a given condition, and represented as a function of time. ImageJ/Fiji plugins were developed to assist for image analysis (semi-automatic segmentation, colocalization and fluorescence intensity analysis, mitochondria tracking and kymograph construction, mitochondria morphology analysis). Those plugins are available upon request.

### Statistical analysis

The proportion and standard deviation of cells positive for mitochondria fluorescence were calculated, and a Chi-squared test was used to compare two groups. Because the chi-squared test assumes independence of experimental units, we also confirmed our p-values by permutations of the population data and calculated the difference in mean between the two groups (10’000 permutations). From this, a normal distribution was derived and used to calculate the probability to observe a difference equally or more extreme than the experimental difference. In all cases, this additional test, which does not require independence of observations, yielded very similar p-values.

### FCCP, Antimycin A, RA

Inhibition of mitochondria activity was performed using FCCP or Antimycin A (Sigma-Aldrich). Retinas were dissected and electroporated with reporter plasmids. Retinas were cultured for 12h, then 5 μM FCCP or 36 μM Antimycin were added to culture medium for 12h. After washing with fresh medium, retinas were kept in culture for additional 12h. Retinas treated with all-trans retinoic acid (Tocris) were electroporated with reporter plasmids and were kept for 36h in culture with 25 nM, 250 nM or 2.5 μM retinoic acid. Fluorescence was analyzed by confocal imaging on fixed retinas. For time-lapse imaging experiments using RA, 2.5 μM retinoic acid in culture medium was added on top of collagen 24h before imaging. Medium was replaced by fresh medium with 2.5 μM retinoic acid at beginning of live imaging session.

### Visualization of mitochondrial membrane potential with TMRM

Retinas were electroporated with reporter plasmids and kept in culture for 8 to 24 h. Retinas were dissociated and plated on chamber slides. TMRM staining was performed using 1 nM TMRM for 20-30 minutes. Following staining, fluorescence was observed on live cells maintained at 37°C under confocal microscope. To assess effect of FCCP on MitoTackerGreen and TMRM signal, retinas were dissociated and stained with 150 nM MTG and 1 nM TMRM for 30 min. After washing, fluorescence was observed on live cells under a Leica Sp5 confocal microscope. 5 μM FCCP was added and fluorescence as recorded every 5 minutes and quantified in imageJ/Fiji.

### RNA-sequencing

Chick E5 retinas were electroporated with Hes5.3-GFP and cultured 8h in medium containing DMSO or 2.5 μM RA. RNA was isolated in triplicate from FACS-sorted Hes5.3-GFP+ cells, and processed for RNA-Seq. Total RNA was quantified with a Qubit (fluorimeter from Life Technologies) and RNA integrity assessed with a Bioanalyzer (Agilent Technologies). The MARTer™ Ultra Low RNA kit from Clontech was used for the reverse transcription and cDNA amplification according to manufacturer’s specifications, starting with 1 ng of total RNA as input. 200 pg of cDNA were used for library preparation using the Nextera XT kit from Illumina. Library molarity and quality was assessed with the Qubit and Tapestation using a DNA High sensitivity chip (Agilent Technologies). Libraries were pooled and loaded at 2 nM for clustering on a Single-read Illumina Flow cell. Reads of 100 bases were generated using the TruSeq SBS chemistry on an Illumina HiSeq 4000 sequencer. The runs generated 56-68 million reads per sample, of which 84.39±1.33% were mappable to reference genome. Mapping raw reads to reference genome was performed using STAR aligner v.2.5.3a to the UCSC Gallus gallus Galgal5 reference. The table of counts with the number of reads mapping to each gene feature of the UCSC Gallus gallus Galgal5 reference was prepared with HTSeq v0.6p1. The differential expression analysis was performed with the statistical analysis R/Bioconductor package edgeR v.3.18.1. Briefly, the counts were normalized according to the library size and filtered. The genes having a count above 1 count per million reads (cpm) in at least 3 samples were kept for the analysis. The raw gene number of the set is 6’789. The poorly or not expressed genes were filtered out. The filtered data set consists of 4’752 genes. The differentially expressed genes tests were done with paired data GLM (general linearized model) using a negative binomial distribution.

### Affymetrix analysis

The 36h microarray transcription profiling described in Chiodini et al., 2013 was designed to test the effects of *Hes5.3* siRNAs, *Atoh7* siRNAs and non-targeting (nt) siRNAs on gene expression at the periphery of the expanding Hes5.3 domain. Briefly, HH24 chick retinas were electroporated with *Hes5.3* siRNAs, *Atoh7* siRNAs or nt siRNA and a *HES5.3-RFP* reporter plasmid. Retina fragments encompassing the peripheral fluorescent domain were micro-dissected under a stereoscopic microscope 36 hours later. RNA was isolated from fragments dissected from four retinas and processed for gene chip analysis. For the 16h microarray analysis, the same siRNAs were used and a similar experimental design was followed. Collection of the retina fragments for RNA isolation was performed 16 hours after electroporation. 150 ng of total RNA were used as input for cRNA preparation using the Ambion WT Expression kit from Ambion (Thermofisher Scientific). After cRNA reverse transcription, single-strand cDNA were fragmented and labeled with the Affymetrix GeneChip WT Terminal Labeling Kit. Targets were hybridized on the GeneChip Chicken Gene 1.0 ST Array. Arrays were washed and stained according to Affymetrix recommendations. GeneChips were scanned on the GS300 Affymetrix scanner. The data were analyzed with Affymetrix Expression Console v1.4.1.46 using Affymetrix default analysis settings and RMA as normalization method. The differential expression analysis was done with a one-way unpaired Anova with the Affymetrix transcriptome Analysis Console v3.1.0.5.

## ACKNOWLEDGEMENTS

We are grateful to Florence Chiodini for valuable help, D. Henrique for the Hes5.3 expression vector, P. Delaunay for the supply of pigeon eggs, J. M. Nunes for his help with statistical analysis, and J.-C. Martinou for inspiring discussions and critical reading of the manuscript. Time-lapse, confocal and electron microscopies were performed at the Bioimaging Platform of the Faculty of Sciences (http://bioimaging.unige.ch). qPCR, RT-qPCR and RNA-Seq experiments were performed at the iGE3 genomics platform of the University of Geneva (http://www.ige3.unige.ch/genomics-platform.php). FACS were performed at the flow cytometry facility of the University of Geneva (https://www.unige.ch/medecine/cytometrie).

## COMPETING INTERESTS

No competing interests declared.

## FUNDING

The Swiss National Science Foundation (grant 31003A-149458), the Gelbert Foundation and the state of Geneva support our laboratory.

## DATA AVAILABILITY

Data from RNA sequencing are available at Gene Expression Omnibus (GEO) with accession number GSE146411. Affymetrix data at 16h are available at GEO with accession number GSE147258. Various scripts used for data processing and analysis can be found at GitHub (https://github.com/lbrodier87).

## AUTHORS CONTRIBUTION

L.B. carried out experiments presented in all Figures except Figure 5 panels B, C, and E, and figure S6 panel C. T.R. brought her expertise in the design of experiments with the pigeon retina, and carried out experiments presented in Figure 5 panels B, C and E, and figure S6 panel C L.B., L.M.S. and J.-M.M. processed and analyzed the data. L.B. and J.-M.M. conceived the study. L.B., L.M.S. and J.-M.M. wrote the manuscript.

## SUPPLEMENTARY MATERIAL

**Figure S1:**
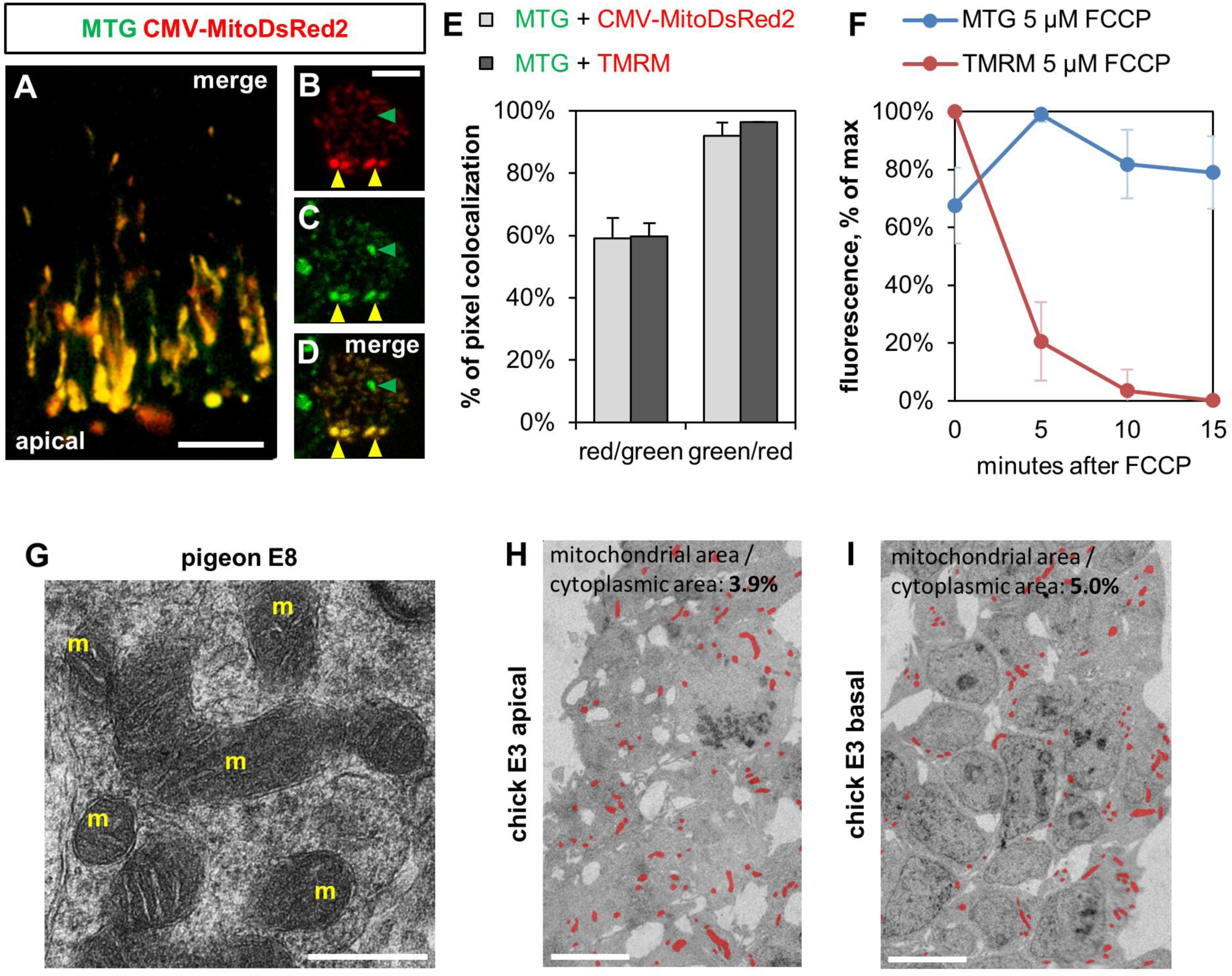
mitochondria detection. (A-D) Co-localization of MitoDsRed2 and MitoTracker Green FM (MTG) labeling. E5 retinas were electroporated with a CMV-MitoDsRed2 plasmid. (A) 24 h later, the retina was incubated with MTG and processed for confocal imaging. (B-D) Red cells were selected by FACS, incubated with MTG and processed for confocal imaging 24 h later. Colocalization of Mito-DsRed2 and MTG in a single cell. Yellow arrowheads: double-labeled mitochondria. Green arrowhead: mitochondria unlabeled with MitoDsRed2. (E) Colocalization of MTG with MitoDsRed2 or TMRM. Light gray bar: E5 retina was electroporated with CMV-MitoDsRed2, cells were dissociated 8h later and stained with MTG. Dark grey bars: dissociated E5 retinal cells were stained with MTG and TMRM. Quantification of the proportion ± s.d. of co-localized pixel for each condition shows that while almost all mitochondria positive for TMRM or MitoDsRed2 are also MTG positive (green/red), lower fractions of MTG positive mitochondria are TMRM or CMV-MitoDsRed2 positive (red/green). (F) Addition of FCCP decreases TMRM signal but has no effect on MTG signal. Dissociated E5 retinal cells were stained with MTG and TMRM. Cells were imaged under confocal microscope at 37°C at 5 minutes interval following addition of μM FCCP Graph shows mean ± s d. of the ratio of fluorescence at each time point relative to maximal fluorescence for 44 cells. (G) TEM showing mitochondria (m) in E8 pigeon retina. (H-I) TEM on apico-basal cross sections of chick retina at E3 (HH20). Mitochondria are artificially colored in red. The ratios of mitochondrial to cytoplasmic areas are similar on the basal and apical sides scale bars: 20 μm (A), 5 μm (B, H, I), 0.05 μm (G)

**Figure S2:**
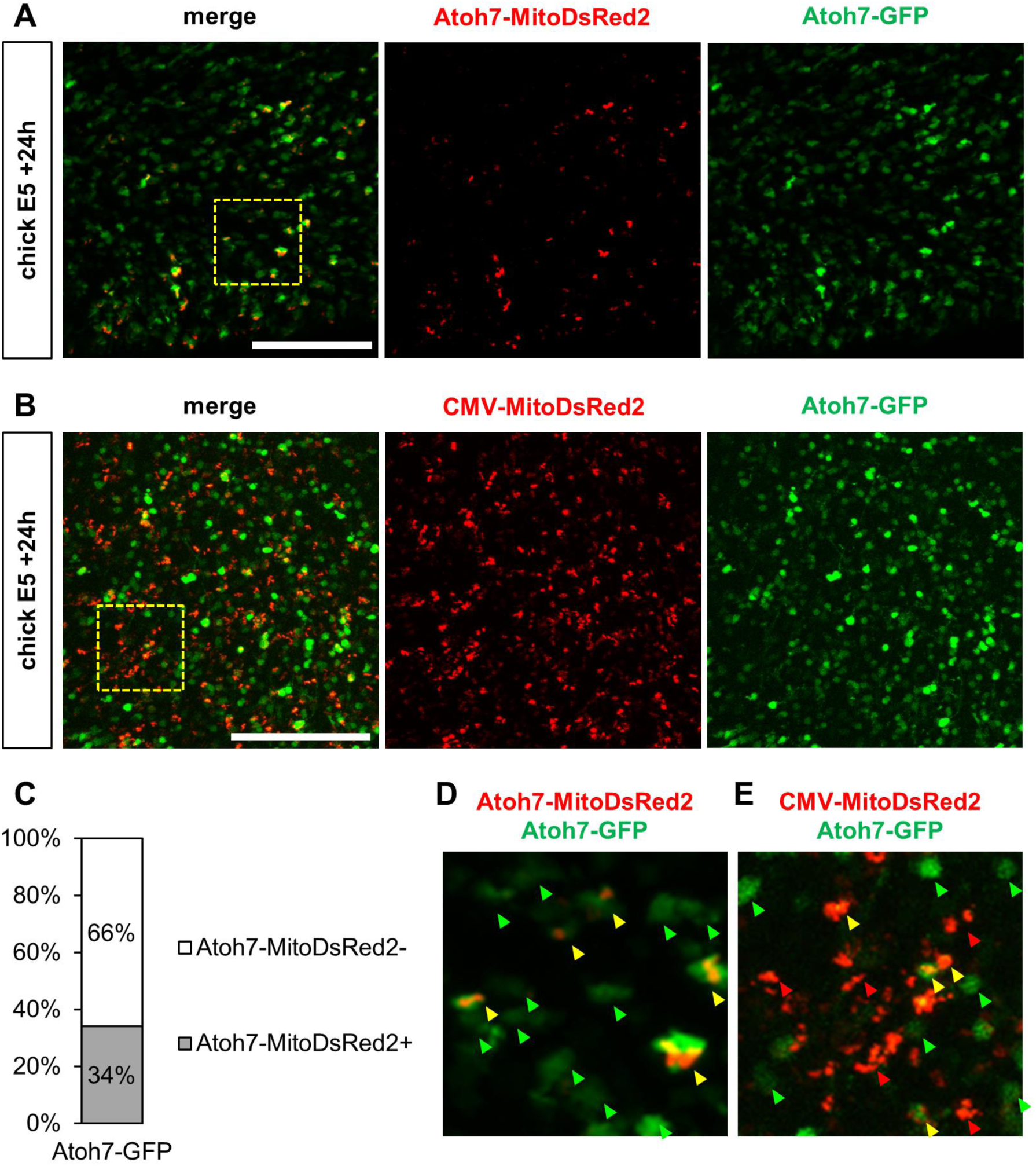
Atoh7-MitoDsRed2 labels mitochondria in a subset of Atoh7-GFP cells. (A-B) Maximal projections from confocal of E5 chick retina co-electroporated with Atoh7-GFP and Atoh7-MitoDsRed2 (A) or with Atoh7-GFP and CMV-MitoDsRed2 plasmids (B) and fixed 24 h later. (C) Mean percentage of GFP+ cells with mitochondria in retinas electroporated with Atoh7-GFP and Atoh7-MitoDsRed2 as displayed in panel A (n=839). (D) Higher magnification of the yellow square from panel A. Fluorescent mitochondria in retinas co-electroporated with Atoh7-MitoDsRed2 and Atoh7-GFP were exclusively located in GFP+ cells. (E) Higher magnification of the yellow square from panel B. The CMV-MitoDsRed2 reporter plasmid does not exhibit this specificity. Green arrowheads: cells positive for Atoh7-GFP and negative for MitoDsRed2 signal; Red arrowheads: cells negative for Atoh7-GFP and positive for MitoDsRed2 signal. Yellow arrowheads: cells positive for Atoh7-GFP and MitoDsRed2 signal.

**Figure S3:**
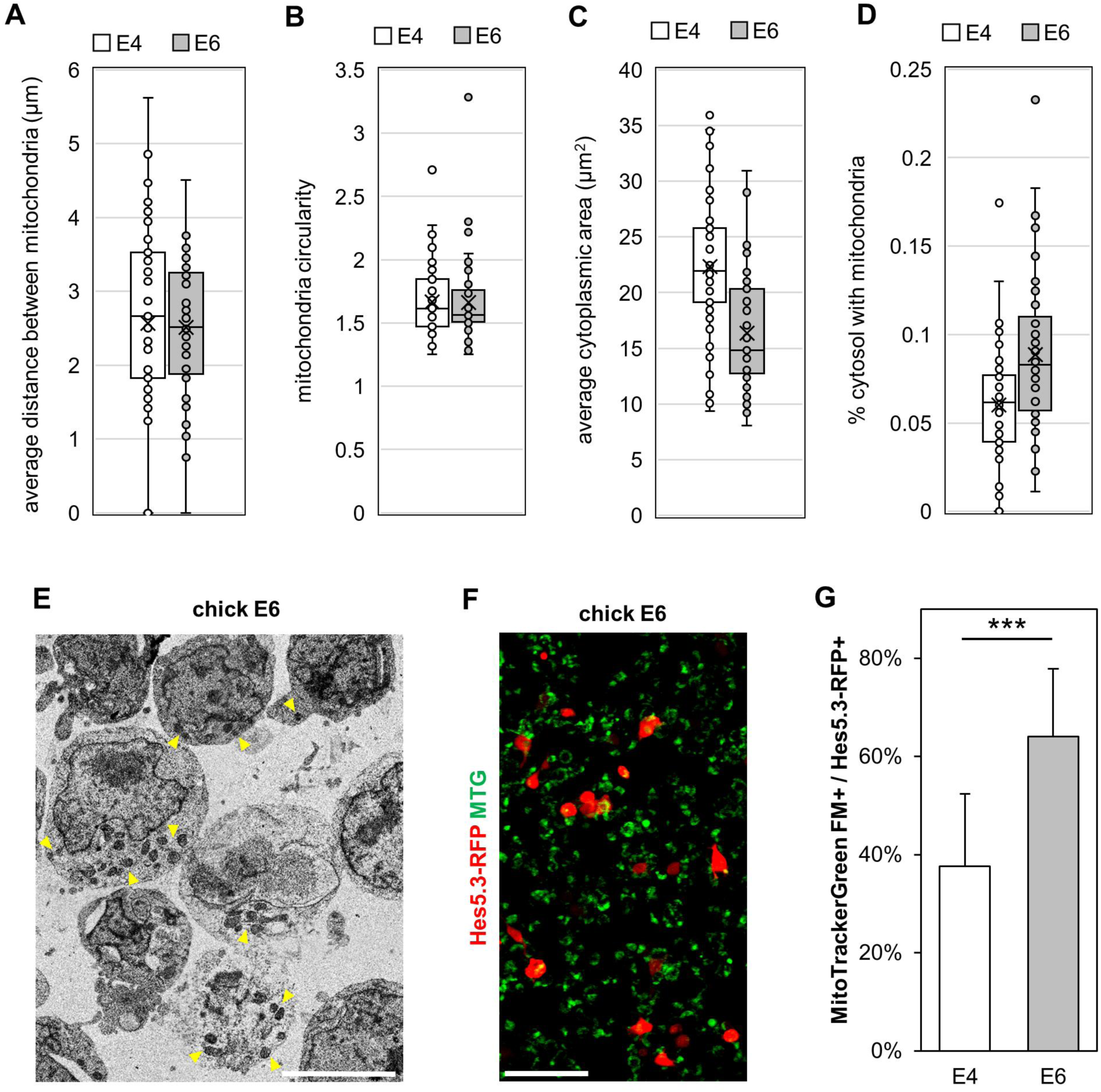
mitochondria in Hes5.3^+^ progenitors. (A-D) Morphometric measurements of mitochondria in individual Hes5.3-GFP+ progenitors (E4 n=51 cells, E6 n=55). (A) Average distance between mitochondria in µm, measured as the distance from centers of fitted ellipses (p<0.05; unpaired t-test; n=51-55). (B) Mitochondria circularity, measured as the ratio of major to minor lengths of fitted ellipse, remains constant (p>0.05; unpaired t-test; n=51-55). (C) The average cytoplasmic area is significantly lower at E6 than at E4 (p<0.001; unpaired t-test; n=51-55 cells). (D) Difference in A is consistent with the increased proportion of cytoplasm filled with mitochondria at E6 (p<0.001; unpaired t-test; n=51-55 cells). (E) An example of a TEM image processed for quantifications shown in A-D and Fig. 3D. Yellow arrowheads point to regions enriched with mitochondria. (F, G) E4 and E6 retinas (4 and 2 respectively, pooled and processed as one biological replicate) were electroporated with Hes5.3-RFP and stained with 150 nM MTG 24 h later. (F) An example of confocal image used for the quantifications shown in G. (G) Average MTG signal relative to Hes5.3-RFP signal measured in multiple images of similar dimensions. MTG density is higher at E6 than at E4 (***p<0.001, unpaired t-test, E4 n=11 images, E6 n=10). Scale bars: 5 μm (E), 50 μM (F)

**Figure S4:**
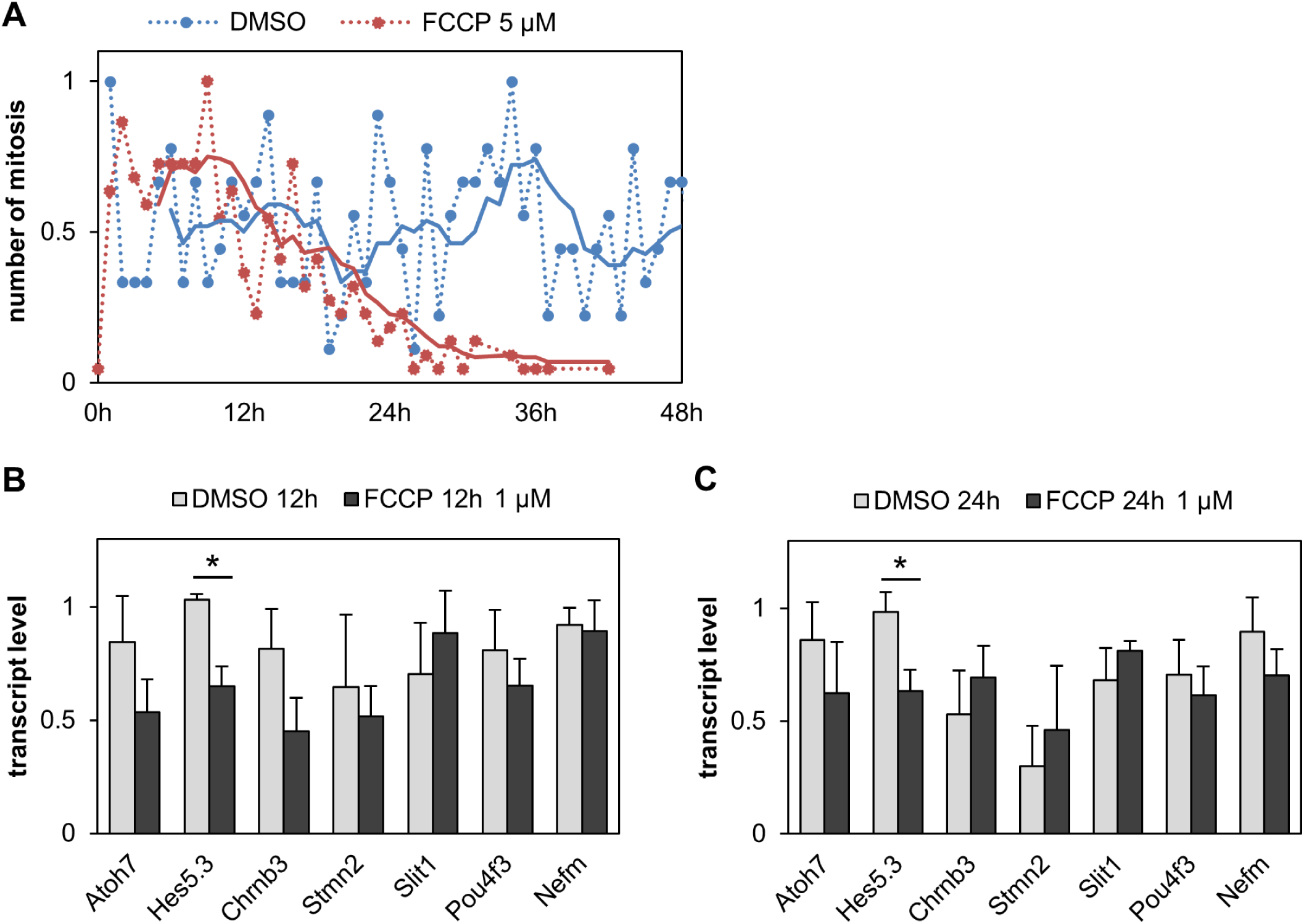
Effect of FCCP on the cell cycle length and expression of RGC markers. Results presented here complement the data provided in Figure 4. (A) E4 retinas were electroporated with Chrna7-GFP and incubated with 5 μM FCCP or DMSO. Plot showing the average mitosis frequency as a function of time in 9 (DMSO) or 22 (FCCP) live imaging movies. (B, C) The right or left retinas from two embryos at E4 were incubated with 1 μM FCCP or DMSO and processed for RT-qPCR analysis 12 h (B) or 24 h (C) later.

**Figure S5:**
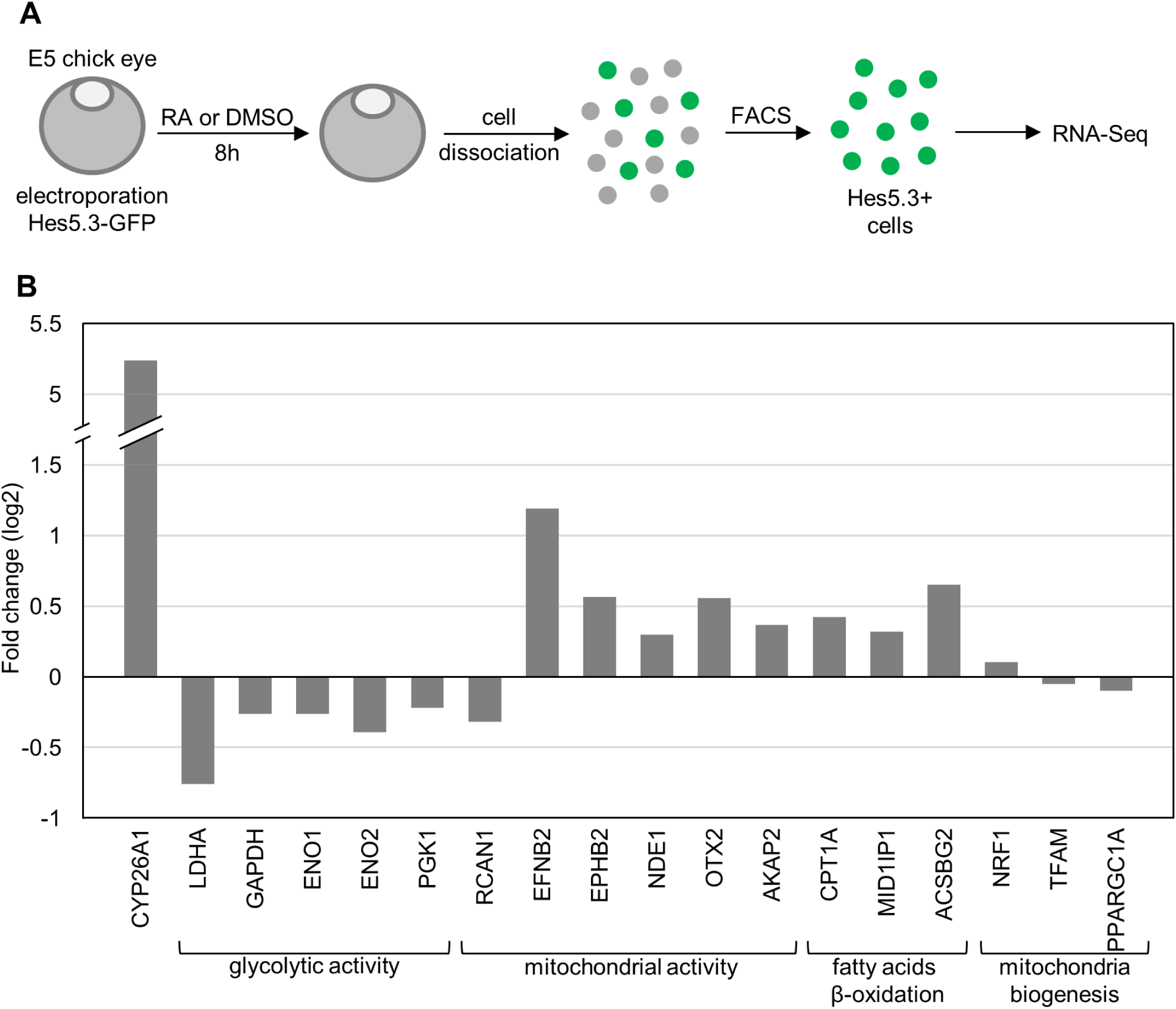
RNA-Seq transcriptome of Hes5.3+ cells. RNA-Seq analysis revealed inhibitory effect of RA on glycolysis in Hes5.3+ pre-committed progenitors. The right or left retinas of 12 embryos at E5 were electroporated with Hes5.3-GFP and cultured in presence of, respectively, 2.3 μM RA or DM for 8 h Retinal cells were dissociated, Hes5.3+ fluorescent cells were isolated by FACS, RNA was extracted and processed for RNA-Seq in triplicate. (A) Schematic representation of the protocol. (B) Bar graph generated from RNA-Seq data showing a selection of genes regulated by RA. Positive or negative fold changes in log2 are presented relative to control DMSO. RA treatment induced an increase of RA-degradative enzyme Cyp26a1, a decrease of expression of five key glycolytic enzymes and of RCAN1, i.e., an inhibitor of mitochondrial metabolism, an upregulation of potential activators of mitochondria metabolism and of fatty acids β-oxidation in mitochondria. Master regulators of mitochondria biogenesis remained unchanged. Statistical analysis: fold changes have p<0.05 except for NRF, TFAM and PPARGC1A.

**Figure S6:**
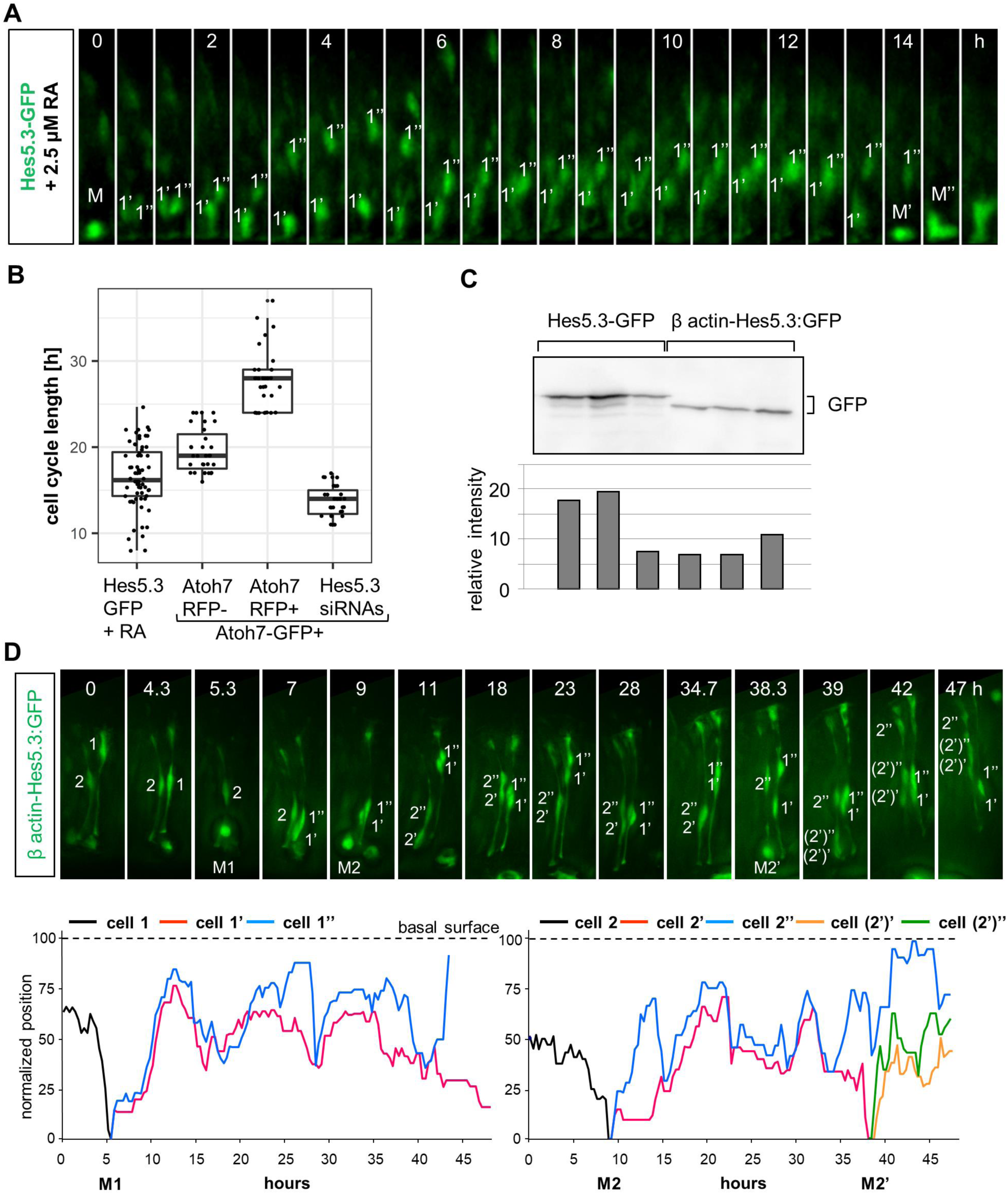
Effects of RA and of HES5.3 on the cell cycle length. (A-B) RA influences the cell cycle length. E4 chick retinas were electroporated with Hes5.3-GFP, 2 μM RA as added and retinas were processed for Time-Lapse imaging 24 h later. (A) Stills from a Time-Lapse movie with a 20 min interval presented as max projections. Hes5.3-GFP+ pre-committed progenitor cells going from mitosis (M) to mitosis (M’ and M’’) he trajectories of the two daughter cells 1’, 1” are shown (B) 64 cells were tracked from mitosis to mitosis. The Atoh7-GFP+ RFP-, Atoh7-GFP+ RFP+ and Hes5.3 siRNAs data are from Chiodini et al. (2013). Hes5.3-GFP+ pre-committed progenitors incubated with RA display cell cycle lengths significantly different from all other groups (p-value < 0.001; unpaired t-test; Hes5.3-GFP + RA n = 64, all other groups n = 31). (C) Similar activities of the *Hes5.3* and β-actin promoters. E4 retinas were electroporated with either a Hes5.3-GFP reporter plasmid or a β-actin-Hes5.3:GFP expression vector. Cytosolic fractions from six retinas were isolated 24 h later and GFP was quantified by western blot. GFP accumulated at similar levels, indicating that *Hes5.3* put under the control of a β-actin promoter is expressed within physiological range. (D) Overexpression of Hes5.3 lengthens the cell cycle. E4 retinas were electroporated with a β actin-Hes5.3:GFP expression vector and fluorescent cells were monitored in real-time 12 h later. (D, upper panel) Stills from a movie spanning 47 h. Positions and mitosis (M) of cell 1 and cell 2 and of their progeny are shown. (D, lower panel) High resolution interkinetic nuclear migration (INM) of cell and 2 and of their progeny 1’, 1” (left) and 2’, 2’’, (2’)’, (2’)’’ (right) Note that only the cell 2’ completes a full cell cycle during the monitoring period.

**Movie S1**

Time-lapse at 20 minutes interval of chick E5 equatorial retina electroporated Atoh7-GFP and Atoh7-MitoDsRed2, showing mitochondria distribution during penultimate and ultimate mitosis of a committed RGC. Mitosis (M), parent cell (yellow arrowhead), daughter cells following first division (red and cyan arrowheads), and daughter cells from 2^nd^ division (blue, orange and green arrowheads) are shown. Time shown as hours:minutes.

**Movie S2**

Time lapse of a RGC axon growing on the basal surface at 22 seconds interval, from a chick E6 retina electroporated with CMV-GFP and CMV-MitoDsRed2. Top: GFP channel, bottom: DsRed2 channel, middle: merge. Growth cone labelled with yellow arrowhead. Time shown as hours:minutes:seconds.

**Table S1.**
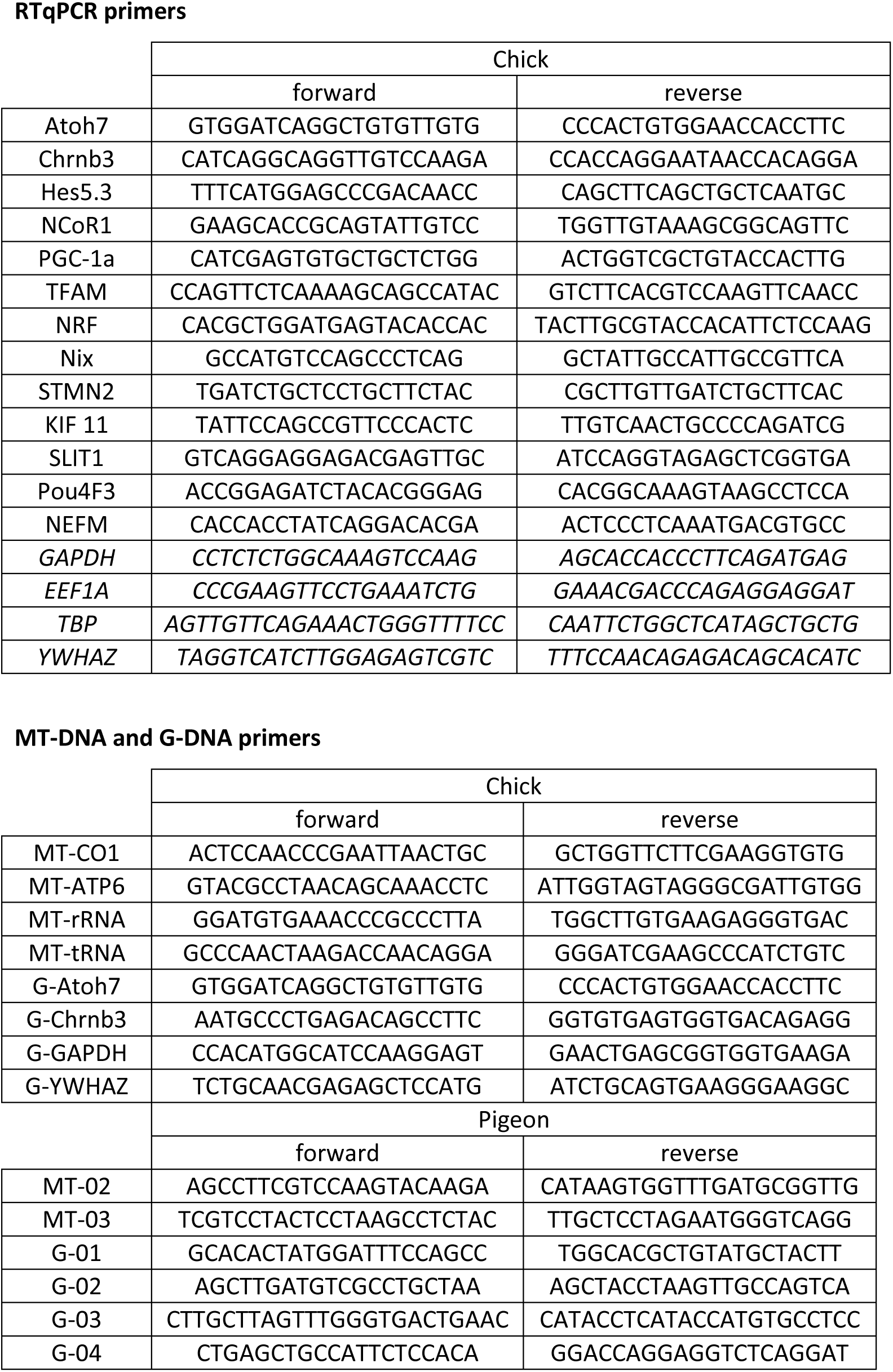
List of primers used in this study for qPCR (mt-DNA, gDNA) and for RT-qPCR (transcripts). Genes used for normalization of RT-qPCR are in italic.

